# The IFT81-IFT74 complex enhances GTP hydrolysis to inactivate RabL2 during early steps of intraflagellar transport

**DOI:** 10.1101/2022.05.31.494111

**Authors:** Niels Boegholm, Narcis A. Petriman, Marta Loureiro Lopez, Tomoharu Kanie, Peter K. Jackson, Jens S. Andersen, Esben Lorentzen

## Abstract

Cilia are important organelles for signaling and motility and are constructed via intraflagellar transport (IFT). RabL2 is a small Rab-like GTPase that localizes to the basal body of cilia via an interaction with the centriolar protein CEP19 before downstream association with the IFT machinery to regulate the initiation of IFT. We have mapped the interaction with RabL2 to residues 107-195 of CEP19, purified the RabL2-CEP19 complex to show that CEP19 is not a GTPase activator protein for RabL2. In contrast, a reconstituted pentameric IFT complex containing IFT81/74 enhances the GTP hydrolysis in RabL2 by 20-fold. The binding site on IFT81/74 that promotes GTP hydrolysis in RabL2 is mapped to a 70 amino acid long coiled-coil region of IFT81/74. We present structural models for minimal IFT81/74-RabL2 complexes and demonstrate that the *Chlamydomonas* IFT81/74 complex enhances GTP hydrolysis of human RabL2 suggesting an ancient evolutionarily conserved function. Our results provide a mechanistic understanding of RabL2 function in the initiation step of IFT and a molecular rationale for why RabL2 dissociates from anterograde IFT trains soon after departure from the ciliary base.

## Introduction

Cilia are slender organelles found on the surface of cells where they serve important functions in motility, sensory reception, and signalling (Rosenbaum and Witman, 2002). Cilia are believed to be ancient organelles present on the last eukaryotic common ancestor and are conserved from unicellular organisms such as the green algae *Chlamydomonas reinhartii* (Cr), a key model organism for ciliary studies, to humans (Dutcher, 2014). Cilium formation is a multi-step process that involves docking of a centriole at the plasma membrane (Sorokin, 1962), vesicular transport from the Golgi to the base of the cilium (Knödler *et al*., 2010; Vetter *et al*., 2015a; Quidwai *et al*., 2021), and construction of the ciliary axoneme (Avasthi and Marshall, 2013). The elongation of the ciliary axoneme requires intraflagellar transport (IFT), the bi-directional trafficking of large proteinaceous particles along the axonemal microtubules to deliver cargo for ciliary assembly (Kozminski *et al*., 1993; Kozminski, Beech and Rosenbaum, 1995; Pedersen and Rosenbaum, 2008). IFT is dependent on kinesin and dynein molecular motors as well as the large multi-subunit IFT complex that mediates the interaction with ciliary cargoes (Kozminski, Beech and Rosenbaum, 1995; Hou, Pazour and Witman, 2004; Bhogaraju, Engel and Lorentzen, 2013; Taschner and Lorentzen, 2016a).

IFT complexes organize into IFT-A and IFT-B sub-complexes that accumulate at the ciliary base (Cole *et al*., 1998; Deane *et al*., 2001). The IFT-B complex can be further sub-divided into a 10 subunit IFT-B1 and a 6 subunit IFT-B2 complex (Taschner *et al*., 2016). IFT-A and IFT-B complexes polymerize into linear assemblies known as IFT trains that are sandwiched between the ciliary axoneme and membrane (Kozminski *et al*., 1993; Kozminski, Beech and Rosenbaum, 1995; Pigino *et al*., 2009). Anterograde IFT trains associate with ciliary cargo such as axonemal components and move from the ciliary base to the tip for cargo delivery (Bhogaraju, Engel and Lorentzen, 2013). Elegant cryo electron tomography (cryo-ET) work has shown that anterograde IFT-B trains organize into 6 nm linear repeat structures whereas the IFT-A trains have a 11 nm repeat (Jordan *et al*., 2018) resulting in an approximate 2:1 ratio for IFT-B:IFT-A complexes in accordance with mass-spectrometry results (Lechtreck *et al*., 2009). At the ciliary tip, the kinesin motor dissociates from the IFT trains and diffuses back to the ciliary base in *Chlamydomonas* (Engel, Ludington and Marshall, 2009). The remaining components of the IFT trains are believed to partly break up before reassembling into dynein driven retrograde IFT trains that have a different ultrastructure (Stepanek and Pigino, 2016; Jordan *et al*., 2018). Except for the kinesin motor, these retrograde trains are thought to consist of the same IFT subunits as anterograde IFT trains (Chien *et al*., 2017). In *C. elegans*, tracking experiments show that IFT-A and IFT-B components have different dwelling times at the ciliary tip, suggesting that IFT-trains are broken into separate IFT complexes (Mijalkovic *et al*., 2018). However, recent work on *Chlamydomonas* show that IFT-A, IFT-B and IFT dynein subcomplexes stay associated through the switch from anterograde to retrograde IFT at the ciliary tip (Wingfield *et al*., 2021).

How do IFT proteins and complexes accumulate at the ciliary base for the initiation of anterograde IFT? Several studies using photo-bleaching of fluorescently tagged IFT subunits have addressed the mechanisms of IFT protein delivery to the base of the cilium. In *Trypanosomes*, experiments with GFP-tagged IFT52 suggested that most IFT material at the ciliary base originates from recycled IFT trains with only a smaller part coming from the cytoplasm (Buisson *et al*., 2013). However, studies of IFT protein dynamics in vertebrate multiciliated cells show that IFT subcomplexes are preassembled in the cytoplasm and recruited to the ciliary base through a diffusion-to-capture mechanism (Hibbard *et al*., 2021). This result agrees with the observation that IFT46 depends on an interaction with IFT52, both subunits of the IFT-B1 complex, for basal body localization in *Chlamydomonas* (Lv *et al*., 2017). A comprehensive study in *Chlamydomonas* uncovered that whereas IFT-A and motor proteins are recruited to the ciliary base from the cytoplasm, IFT-B proteins are both recruited from the cytoplasm as well as from ‘re-used’ retrograde IFT trains (Wingfield *et al*., 2017). Anterograde IFT cargo such as tubulin and IFT dynein are loaded onto anterograde IFT trains shortly before departure (Wingfield *et al*., 2017).

The mechanism of IFT train assembly at the ciliary base was also addressed by cryo-ET in a recent seminal study demonstrating that IFT trains assemble in a sequential manner at the base of the cilium (Hoek *et al*., 2021a). IFT train assembly appears to occur first through polymerization of IFT-B followed by IFT-A polymerization and lastly association of IFT motors (Hoek *et al*., 2021). Photobleaching experiments in *Chlamydomonas* show that IFT and motor proteins recover at different rates (3-10s) with IFT43, IFT20 and IFT54 requiring about 9s for full recovery (Wingfield *et al*., 2017). This result suggest that the timescale of IFT train assembly at the ciliary base is in the order of seconds, and is followed by injection into the cilium via an avalanche-like mechanism (Ludington *et al*., 2013; Wingfield *et al*., 2017; Hoek *et al*., 2021).

Although the process of IFT initiation at the base of the cilium is not well understood, several lines of evidence suggest that the IFT-B complex plays a crucial role. The IFT-B polymers appear to form first and subsequently serve as a scaffold for the remaining IFT train components (Hoek *et al*., 2021). Furthermore, tomographic reconstructions show that the IFT-B complex contacts the kinesin-II motor required for initiating and driving anterograde IFT (Jordan *et al*., 2018), an interaction that likely occurs through the IFT88/52/57/38 hetero-tetramer (Funabashi *et al*., 2018). The IFT-B1 complex contains the two small GTPases IFT22 and IFT27 (Taschner and Lorentzen, 2016b). Small GTPases regulate many cellular processes by cycling between an inactive GDP-bound conformation and an active GTP-bound conformation that interacts with downstream effectors (Wittinghofer and Vetter, 2011). Activation through GDP→GTP exchange is promoted by guanine nucleotide exchange factors (GEFs) whereas inactivation through GTP hydrolysis is promoted by GTPase activating proteins (GAPs). IFT27 (aka Rab-like 4 (RabL4)) associates with IFT25 to form a hetero-dimer (Qin *et al*., 2007; Wang *et al*., 2009; Bhogaraju *et al*., 2011) and was initially suggested to play a role in IFT initiation (Wang *et al*., 2009). However, several subsequent studies have shown that IFT25/27 is dispensable for anterograde IFT but is instead required for the ciliary export of the BBSome complex and associated retrograde cargoes including sonic hedgehog signalling factors in mammals (Keady *et al*., 2012; Eguether *et al*., 2014; Liew *et al*., 2014; Dong *et al*., 2017) and phospholipase D in *Chlamydomonas* (Lechtreck *et al*., 2009, 2013). IFT22 (aka RabL5) was initially discovered in *Chlamydomonas* (Wang *et al*., 2009) where it regulates the cellular levels of IFT proteins (Silva *et al*., 2012). However, IFT22 does not appear to be required for IFT initiation but, together with BBS3, is involved in recruiting the BBSome to the ciliary base (Xue *et al*., 2020). In *Caenorhabditis elegans*, mutation of IFT22 also does not affect ciliogenesis or IFT (Schafer *et al*., 2006; Inglis, Blacque and Leroux, 2009). In contrast, IFT22 in *Trypanosoma brucei* does appear to be required for proper ciliogenesis as IFT22 knockdown results in a retrograde IFT phenotype characterized by short cilia full of IFT material (Adhiambo *et al*., 2009; Wachter *et al*., 2019). However, the retrograde IFT phenotype of IFT22 knockdown cells suggests that IFT22, like IFT27, is not required for IFT initiation.

More recently, a third GTPase, RabL2, was shown to associate with the IFT-B complex and regulate IFT initiation and cilium formation (Kanie *et al*., 2017; Nishijima *et al*., 2017). RabL2 is required for proper ciliogenesis in both *Chlamydomonas* (Nishijima *et al*., 2017) and in mammalian cells (Kanie *et al*., 2017). Mutations in RabL2 cause ciliopathies including male infertility because of defects in the assembly of cilia of sperm cells (Lo *et al*., 2012; Ding *et al*., 2020). Furthermore, RabL2 controls the ciliary localization of G-protein coupled receptors (GPCRs) in primary cilia suggesting a conserved role in the assembly/function of both motile and primary cilia (Dateyama *et al*., 2019). This agrees with the evolutionary conservation of RabL2 in ciliated species and the lack of RabL2 in non-ciliated eukaryotes (Eliáš *et al*., 2016). RabL2 is recruited to the basal body of cilia via an interaction with the centriolar protein CEP19 (Jakobsen *et al*., 2011) and subsequently handed over to the IFT-B complex to initiate IFT at the ciliary base (Kanie *et al*., 2017). Knockout of CEP19 or RabL2 significantly reduces the number of IFT trains in cilia suggesting a crucial function for RabL2 in controlling the injection of IFT trains into cilia (Kanie *et al*., 2017). Wild-type RabL2 was shown to dissociate from IFT trains shortly after departure from the ciliary base whereas a GTP-locked RabL2 variant (Q80L in human RabL2) stays associated with IFT trains and accumulates in cilia (Kanie *et al*., 2017).

In contrast, the S35N RabL2 mutant unable to bind GTP does not rescue ciliogenesis defects of RabL2 knockout cells (Kanie *et al*., 2017). Moreover, a recent study suggests that GTP-hydrolysis of RabL2 is required for export of the BBSome complex and receptor cargoes out of cilia (Duan *et al*., 2021). There is thus ample evidence that the nucleotide state of RabL2 is important in RabL2 regulation of IFT initiation and cilium function.

Here, we present a comprehensive biochemical analysis of RabL2 and the association with CEP19 and the IFT-B1 complex. We show that the IFT complex, rather than CEP19, functions as a GAP that stimulates GTP hydrolysis to inactivate RabL2 and demonstrate that this activity is conserved from *Chlamydomonas* to human. Our data allow us to present a model where RabL2 associates with IFT trains to licence initiation at the ciliary base, followed by stimulation of GTP-hydrolysis, which inactivates RabL2 to trigger its dissociation from IFT trains.

## Results

### CEP19 has strong affinity for RabL2-GTP but is not a GAP for RabL2

RabL2 was previously shown to locate to the basal bodies of cilia via an interaction with the protein CEP19 (Kanie *et al*., 2017). We purified wild-type (WT) and GTP-locked Q83L mutant CrRabL2 (Fig. S1A-D) and demonstrated that RabL2 does not carry over nucleotides during purification (Fig. S1E-F), which is consistent with the reported low micromolar affinity of RabL2 for GTP/GDP nucleotides (Kanie *et al*., 2017). In addition, we purified CrCEP19 as a C-terminal truncation encompassing residues 1-208 (CrCEP19_1-208_, Fig S1G-H). The C-terminal 40 residues of CrCEP19 are predicted to be intrinsically disordered and were thus omitted in the CEP19_1-208_ construct (Fig. S1I-J). The interaction between RabL2 and CEP19 was studied by size-exclusion chromatography (SEC) and isothermal titration calorimetry (ITC). The results show that GTP-bound RabL2 co-purifies with CEP19_1-208_ on SEC at an elution volume that is significantly shifted when compared to CEP19_1-208_ suggesting the formation of a stable complex (Fig. 1A). Furthermore, this result shows that RabL2/CEP19 complex formation only requires the N-terminal 208 residues of CEP19 (Fig. 1A-B). To obtain quantitative data on the affinities between RabL2 and CEP19, GTP- or GDP-bound RabL2 was titrated with CEP19_1-208_ in ITC experiments. The ITC experiments show that the CrRabL2-GTP/ CEP19_1-208_ complex has a dissociation constant (Kd) of 0.28μM (Fig. 1C). Interestingly, GDP-bound RabL2 still associates with CEP19, but with a Kd of 12.7 μM for the CrRabL2-GDP/CEP19_1-208_ complex (Fig. 1D). The affinity of CEP19 for RabL2-GDP is thus 45 times lower than for RabL2-GTP. This result agrees with the notion that CEP19 senses the nucleotide state of RabL2 but is not a true effector for RabL2. Given that CEP19 preferably associates with the GTP-bound state of RabL2, we tested if CEP19 functions as a GAP for RabL2 using GTPase assays with CrRabL2 alone or in complex with CrCEP19_1-208_. The results of the GAP assay show that CEP19 does not have any stimulating effect on the GTP hydrolysis rate of RabL2 demonstrating that CEP19 is not a GAP for RabL2 (Fig. 1E).

**Figure 1:**
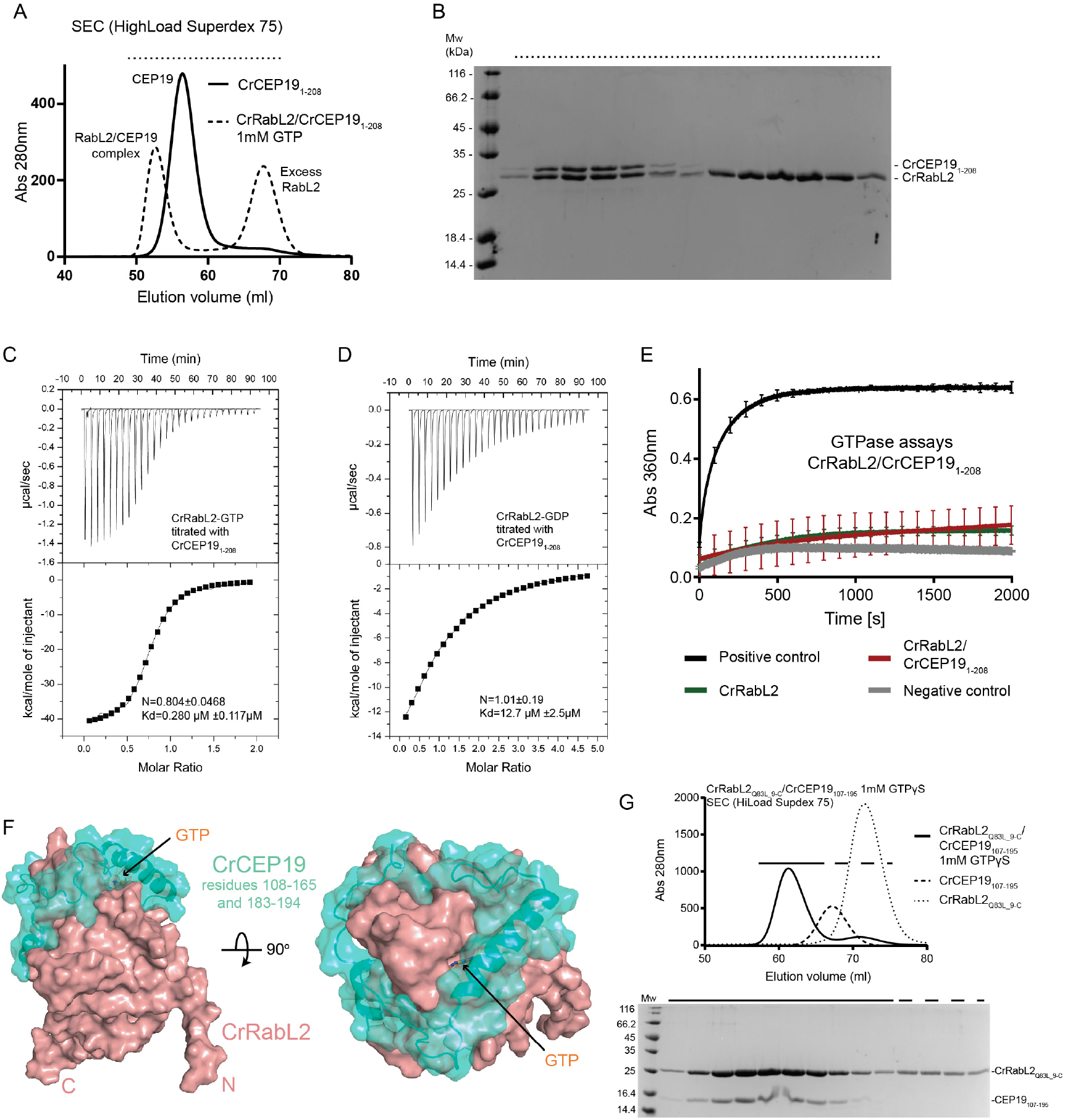
GTP- and GDP-dependent RabL2-CEP19 complex formation. (A) SEC profile that shows the co-purification of CrCEP19_1-208_ and CrRabL2 in the presence of GTP. The elution volume of CrCEP19-RabL2 is significantly shifted compared to the volumes of CEP19 or RabL2 alone demonstrating the formation of a complex **(B)** Coomassie-stained SDS-PAGE of the SEC fractions highlighted in (A) with a horizontal dashed line on top. **(C-D)** ITC measurements of purified CEP19 titrated with CrRabL2 in the presence of GTP (C) or GDP (D). **(E)** GTPase assays of RabL2 alone or in complex with CEP19_1-208_ measuring the release of inorganic phosphate upon GTP hydrolysis as a function of time. Each experiment was carried out in triplicates, curves are averages of these triplicates, and the error bars indicate standard deviation of measurements for every 100s. Inorganic phosphate was used as the positive control and GTP in buffer as the negative control. **(F)** Surface representation of the AlphaFold predicted structure for the complex between CrCEP19_108-194_ (green color) and CrRabL2 (red-salmon color). **(G)** SEC profile (top) and Cooomassie stained SDS gel (bottom) of the complex between RabL2 and a minimal binding-region of CEP19 (residues 107-195) demonstrating a direct physical interaction.

There are currently no experimentally determined structures available for RabL2 or CEP19 proteins. We thus carried out structural modelling of the CrRabL2/CEP19 complex structure using alphafold multimer (Jumper *et al*., 2021; Evans *et al*., 2022) (Fig. 1F and S1I-J). With exception of the N-terminal 20 residues and the C-terminal 40 residues, the structural model for CrRabL2 was predicted with very high confidence (predicted local-distance difference test (pLDDT) score >90) encompassing the entire core GTPase fold (Fig. S1I). CrCEP19 is mostly predicted to fold into 4 α-helices interspaced by long loop regions likely to represent intrinsically disordered regions (Fig. S1I). However, two helices (residues 184-193 and 120-137) and two regions without secondary structure (residues 108-119 and 138-165) of CrCEP19 are predicted with very high confidence and form close contacts with RabL2 (Fig. 1F and S1I). The low predicted aligned error (PAE) between residues in these regions of CrCEP19 and CrRabL2 residues (Fig. S1J) suggests that they are involved in CEP19/RabL2 complex formation (Fig. 1F). These regions of CEP19 are predicted to encircle RabL2 forming a crown-like structure (Fig. 1F). Interestingly, residues 120-137 of CrCEP19 form an α-helix that lines the nucleotide binding pocket of RabL2 and these residues likely sense the nucleotide state of RabL2 thus providing increased affinity for GTP-bound RabL2. The remaining RabL2-interacting parts of CEP19 (residues 140-165 and 183-194) likely provide nucleotide-independent interactions that allow complex formation of CEP19 with RabL2-GDP. To validate the structural model shown in Fig. 1F and confirm the CEP19 minimal binding region for RabL2, CrRabL2/CEP19_107-195_ was reconstituted and co-purified by SEC demonstrating the formation of a stable complex (Fig. 1G). The data presented in Figs. 1 and S1 allow us to conclude that residues 107-195 of CEP19 constitute a minimal binding region that prefers the GTP-bound state of RabL2 but does not stimulate the GTP hydrolysis by RabL2.

### Reconstitution of RabL2-containing IFT-B1 complexes

RabL2 in the GTP-bound state was previously shown to associate with the IFT complex via IFT81/74 (Kanie *et al*., 2017; Nishijima *et al*., 2017). Recently, visual immunoprecipitation experiments with full-length IFT81 and different IFT74 truncations showed that the binding site for RabL2 is located on the IFT74/81 heterodimer N-terminally to the IFT27/25 heterodimer (Zhou *et al*., 2022). We have recombinantly expressed and purified *Chlamydomonas* and human hetero-hexameric IFT-B1 complexes containing RabL2 in the presence of the non-hydrolysable GTP analogue GTPγS (IFT81/74/27/25/22/RabL2, IFT-B1 hexamer, see Fig. 2A-C). In humans, RABL2 is represented by two nearly identical paralogs namely RABL2A and RABL2B (Wong *et al*., 1999). As these are only differentiated by 3 amino acids and are functionally equivalent in the rescue of the ciliogenesis defect of the RABL2A;RABL2B double knockout cells (Kanie *et al*., 2017), we used the RABL2B paralog for reconstitution of IFT-B1 complexes. We were unable to express the human IFT-B1 hexamer in *E.coli* but did succeed in purification using insect cells as a eukaryotic expression system, although the yield obtained was much lower than for the *Chlamydomonas* counterpart using *E.coli* as an expression system.

**Figure 2:**
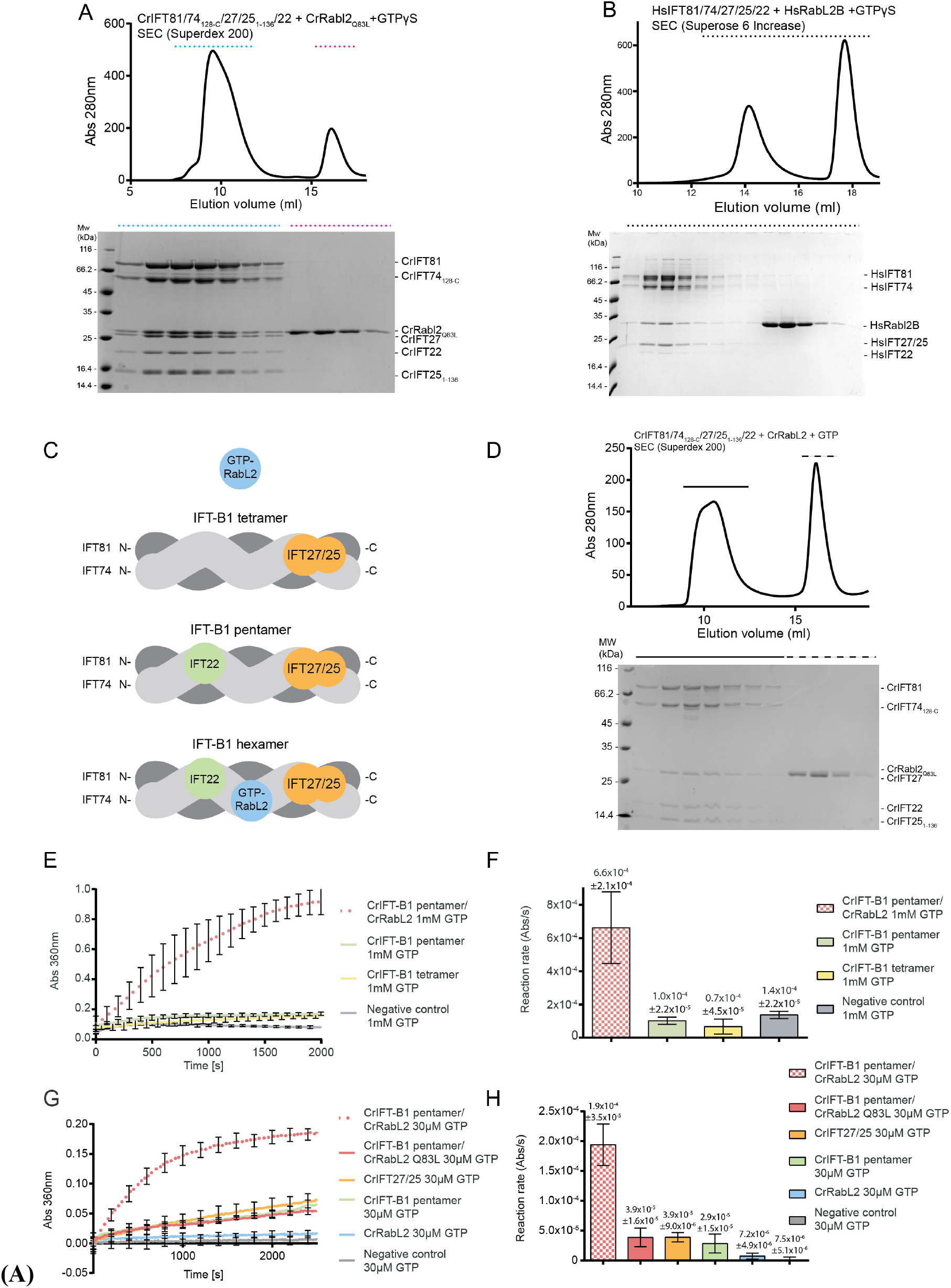
Purification and GTPase activity of IFT-B1 complexes. **(A)** SEC profile of CrRabL2 co-purification with CrIFT81/74/27/25/22 (top) in the presence of non-hydrolysable GTP homologue, GTPγS. Coomassie stained SDS gel of SEC fraction highlighted by dashed lines (bottom). **(B)** SEC profile for the purification of the HsIFT81/74/27/25/22/RabL2 complex in the presence of GTPγS (top) and the corresponding coomassie stained SDS gel (bottom). **(C)** Schematic representation of the IFT-B1 tetramer, pentamer and hexamer. **(D)** SEC profile of the incubation of the CrIFT-B1 pentamer in the presence of CrRabL2 and GTP for 3h at room temperature. The gel at the bottom shows that RabL2 does not stay associated with the IFT-B1 pentamer under these conditions. **(E, G)** GTPases assays with the indicated proteins following the release of inorganic phosphate upon GTP hydrolysis as a function of time. Each experiment was carried out in triplicates. The curves represent averages and error bars indicate standard deviation of measurements for every 100s for panel (E) and every 200s for panel (G). **(F, H)** Quantification of the reaction rates (arbitrary units of Absorbance (Abs) per second(s)) for each experiment shown in panels (E, G) calculates using linear regression of the first 500s and standard deviations are given for 3 independent experiments.

When GTPγS was added, stable hexameric IFT-B1 complexes could be purified by SEC using both WT RabL2 and Q83L mutant of CrRabL2 deficient in GTP hydrolysis (Fig 2A-B). Interestingly, when WT CrRabL2 and GTP was mixed with the IFT-B1 complex and incubated for 3h at room temperature, RabL2 no longer associated with the IFT-B1 complex, perhaps suggesting hydrolysis of GTP over time (Fig. 2D). In the *Chlamydomonas* complex containing IFT74ΔN and IFT25ΔC, all three small GTPases of the IFT-B1 complex (IFT27, IFT22 and RabL2) are present in apparent stoichiometric amounts suggesting that that their association with IFT-B1 is not mutually exclusive and that each GTPase likely has a unique binding site within the IFT-B1 complex (Fig. 2A). The human hexameric IFT-B1 complex contains full length subunits resulting in the co-migration of IFT27 and IFT25 on the SDS gel (Fig. 2B). As shown by Kaine et al., we observe that RabL2 association with the IFT-B complex (IFT81/74/27/25/22, IFT-B1 pentamer, see Fig. 1C) is completely dependent on GTP as RabL2 does not co-purify with IFT-B1 in the absence of a non-hydrolysable GTP analogue such as GTPγS (Fig. S2A). In agreement with previous publications, IFT22 and IFT27 do not require the addition of GTP to associate with the IFT-B1 complex (Fig. S2A) (Taschner *et al*., 2014; Wachter *et al*., 2019). Quantitative data on the affinity of CrRabL2 for the CrIFT-B1 complex were obtained from ITC experiments revealing a Kd of 0.59μM for the RabL2-GTPγS bound IFT-B1 complex (Fig. S2B). No binding was observed between RabL2-GDP and IFT-B1 suggesting that the affinity is at least two orders of magnitudes lower than for GTP-bound RabL2 (Fig. S2C). These experiments verify that the IFT-B complex associates only with the active GTP-bound conformation of RabL2 and indicate GTP-hydrolysis over time resulting in dissociation of RabL2 from the IFT-B1 complex.

### The IFT-B1 complex is a GAP for RabL2 but not for IFT27 or IFT22

Given that IFT81/74 is the binding platform for the 3 small GTPases IFT22, IFT27 and RabL2, we asked if IFT81/74 functions as a GAP for one or more of the small GTPases. In addition to the CrIFT-B1 pentamer and hexamer (Fig. 2A and S2A), we purified the *Chlamydomonas* IFT81/74ΔN/27/25ΔC complex (IFT-B1 tetramer), which contains IFT27 but lacks both RabL2 and IFT22 (Fig. S2D). The IFT-B1 tetramer does not require the addition of GTP for IFT27 to stay associated with the complex. Additionally, we showed in a previous study that IFT81/74/22 complexes co-purify with GTP although GTP is not mandatory for association of IFT22 with the IFT complex (Wachter *et al*., 2019). In contrast, RabL2 alone does not co-purify with GTP (Fig. S1E-F) and requires the addition of GTP to form a complex with IFT81/74 (Fig. 2A-B and S2A) (Kanie *et al*., 2017). Interestingly, while we observed that RabL2 incubated with the IFT-B1 pentamer and GTPγS resulted in co-purification of an IFT-B1 hexamer on SEC (Fig. 2A), the incubation of RabL2 with GTP and the IFT-B1 pentamer result in separate elution peaks for RabL2 and the IFT-B1 pentamer demonstrating that an IFT-B1 hexamer was not formed (Fig. 2D). Given that the intrinsic GTP hydrolysis rate of RabL2 is very low (Fig. 1E), this result could indicate that GTP was hydrolysed during incubation with the IFT-B1 pentamer perhaps suggesting an increased rate of GTP hydrolysis when RabL2 associates with the IFT-B complex.

To analyse the potential GAP function of IFT-B1 complexes, GTPase assays were carried out with purified complexes to measure the GTP hydrolysis rates. Initial GTPase activity assays with the IFT-B1 tetramer, pentamer or hexamer demonstrated that tetrameric and pentameric complexes without RabL2 do not have GTPase activity above background levels when using 1mM GTP in the assay (Fig. 2D). This result suggests that the IFT81/74 is not a GAP for IFT22 or IFT27. In contrast, the hexameric IFT-B1 complex containing RabL2 displayed robust GTP hydrolysis activity with a reaction rate about 5-fold higher than what was observed for IFT complexes lacking RabL2 (Fig. 2D-E). These data indicate that incorporation of RabL2 into the IFT complex activates the GTP hydrolysis in RabL2. Alternatively, RabL2 could act as a GAP towards IFT27 or IFT22 in context of the IFT-B1 complex. To further analyse the GTP hydrolysis activity of the IFT-B1 complex and distinguish between these two possibilities, GTPase assays were repeated using WT or RabL2_Q83L_ catalytic mutant reconstituted IFT-B1 hexamers (Fig. 2F-G). Under the conditions of the assay (single turnover kinetics using 30 μM GTP), WT RabL2 in context of the hexameric IFT-B1 complex has 26-fold higher GTPase activity than WT RabL2 alone (Fig. 2F-G). Adjusting for the low basal GTPase activity of IFT27 and IFT22 within the IFT-B1 hexamer, the IFT-B1 complex increases the reaction rate of RabL2 by approximately 20-fold under the conditions of the assay in Fig. 2G. Several protein families within the superfamily of small GTPases rely on a catalytic glutamine from the switch 2 region for GTP hydrolysis (Pai *et al*., 1990, 1990; Seewald *et al*., 2002). This catalytic glutamine is conserved in most Rab proteins including RabL2 (Q83 in CrRabL2 and Q80 in HsRabL2B) but is not conserved in most IFT27 or IFT22 sequences (Bhogaraju *et al*., 2011). Mutation of this catalytic glutamine to leucine (CrRabL2_Q83L_) is thus expected to abolish GTP hydrolysis in RabL2. Indeed, the hexameric IFT-B1 complex containing the RabL2_Q83L_ mutant has the GTPase activity reduced to background levels observed for the IFT-B1 pentamer without RabL2 (Fig. 2G-H). This result shows that the increase in GTPase activity of hexameric compared to pentameric or tetrameric IFT-B1 complexes is a result of GTP hydrolysis in the active site of RabL2 and confirms that Q83 is important for catalysis. It is noteworthy that the GTPase activity observed for the IFT-B1 pentamer, when compared to background levels, can be recapitulated by the IFT27/25 complex suggesting that the low level of GTP hydrolysis observed for the IFT-B1 pentamer can be attributed to IFT27 rather than IFT22 (Fig. 2G-H). Furthermore, this result verifies that the intrinsic GTPase activity of IFT27/25 is not increased in context of the IFT-B1 complex. We conclude that the IFT-B complex is a GAP for RabL2. Association of GTP-bound RabL2 with the IFT-B complex will thus lead to increased GTP hydrolysis to inactivate RabL2 resulting in the subsequent dissociation of GDP-bound RabL2 from the IFT-B complex.

### A minimal IFT81_460-533_/74_460-532_ complex binds RabL2 and stimulates GTPase activity

To biochemically map the binding site for RabL2 on IFT81/74, we reconstituted and purified *Chlamydomonas* complexes harbouring truncated IFT81 and IFT74 proteins. Removing the most N-terminal 150 residues of both IFT81 and IFT74 did not impact the ability of IFT27/25 or RabL2 to co-purify with IFT81/74 in a complex on SEC confirming that the N-termini of IFT81 or IFT74 are not required for complex formation with RabL2 or IFT27/25 (Fig. S3A). However, deleting the 150 C-terminal residues (IFT81_132-475_/74_132-475_) disrupts binding of RabL2 while retaining the ability to associate with IFT22 (Fig. S3B). Importantly, a minimal complex containing the last 3 coiled-coil segments of IFT81_460-C_/74_460-C_ retains the ability to associate with both RabL2 and IFT27/25 (Fig. S3C). Finally, we show that the most C-terminal residues following the last coiled coil segment of IFT81/74 are not required for binding of RabL2 or IFT27/25 (Fig. S3D). These experiments biochemically map the binding region for CrRabL2 to a coiled-coil segment between residues 460-623 of CrIFT81 and residues 460-615 of CrIFT74. Further trimming of the C-termini of IFT81 and IFT74 resulted in a predicted coiled-coil segment of about 70 residues (IFT81_460-533_/74_460-532_) that co-purifies with RabL2-GTPγS to yield a stable complex on SEC (Fig. 3A). Furthermore, pull-down experiments with RabL2 demonstrated that IFT81_460-533_/74_460-532_ is sufficient to recapitulate RabL2 binding in the presence of GTPγS but not in the presence of GDP (Fig. 3B-C). These results show that a short 70 residue coiled-coil fragment of IFT81/74 constitutes a GTP-dependent minimal RabL2 binding region. The fact that IFT81_460-533_/74_460-532_ discriminates between GTP- and GDP-bound RabL2 conformations suggests that complex formation involves the GTPase switch regions of RabL2.

**Figure 3:**
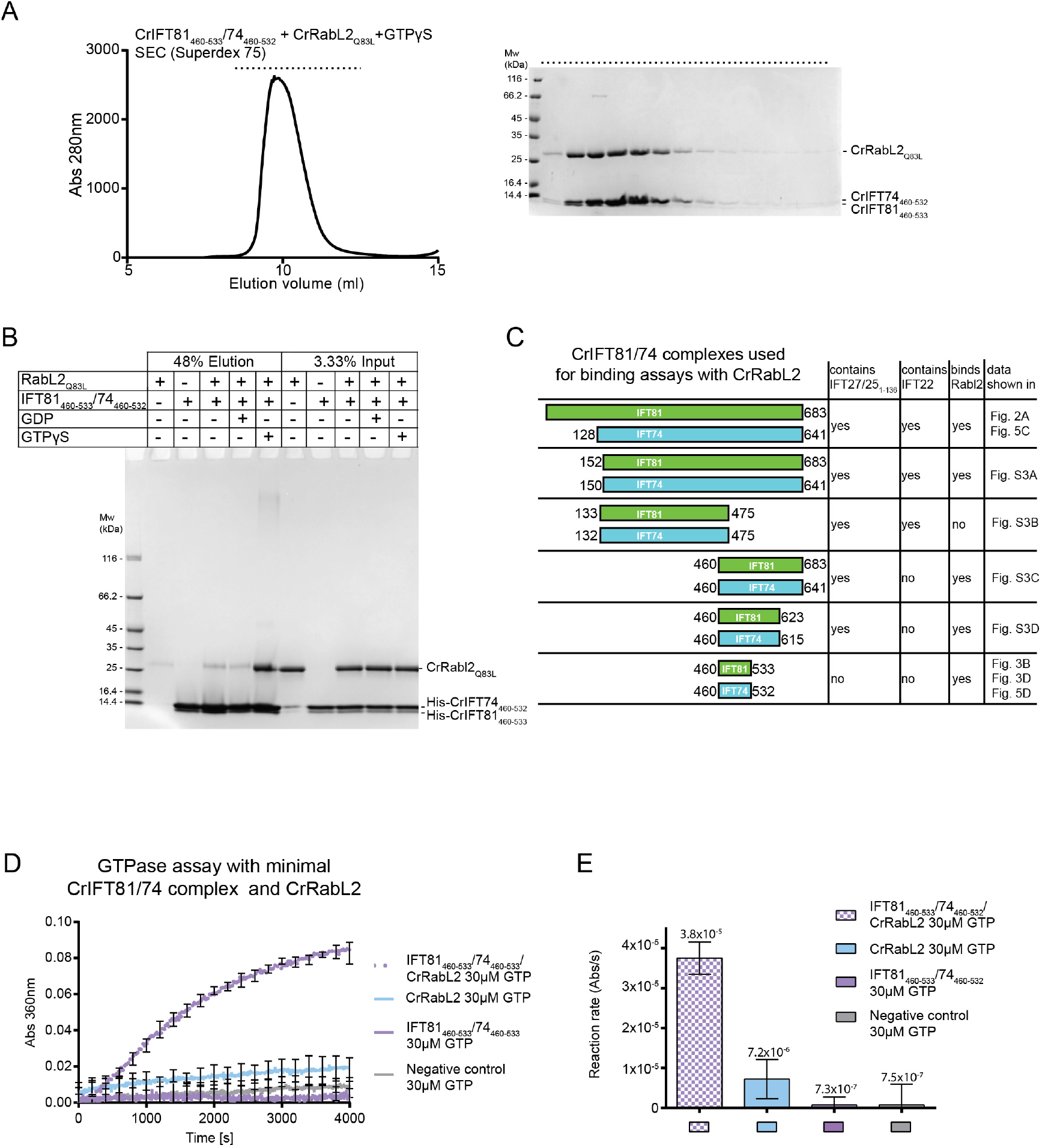
mapping of a minimal CrIFT81_460-533_/CrIFT74_460-532_ complex that binds RabL2 and activates GTP hydrolysis. **(A)** SEC profile showing that a minimal CrIFT81_460-533_/CrIFT74_460-532_ complex co-purifies with CrRabL2 in the presence of GTPγS (left). The right panel displays the Coomassie stained SDS gel of SEC fractions (horizontal top dashed line). **(B)** An N-terminal hexa-histidine tagged CrIFT81_460-533_/74_460-532_ complex interacts with untagged CrRabL2 in a GTPγS dependent manner in pulldown assays. **(C)** Schematic of all CrIFT81/74 truncations used in this study for Rabl2 binding assays. The presence of CrIFT27/25_1-136_ and/or CrIFT22 in a complex with the CrIFT81/74 variants as well as their ability to bind CrRabl2 is indicated. The corresponding data are found as indicated in the last column. **(D)** GTPase using CrRabL2 and a minimal IFT81/74 complex show that this is sufficient to increase GTP hydrolysis by RabL2. Each experiment was done in triplicates, curves represent the averages with error bars representing standard deviations each 200s. **(E)** Quantification of GTPase reaction rates (arbitrary units of Absorbance (Abs) per second(s)) using linear regression of the curves in panel (D). Error bars indicate standard deviation for three independent experiments.

To test if IFT81_460-533_/74_460-532_ is sufficient to stimulate the GTPase activity of RabL2, GTPase assays were carried out with IFT81_460-533_/74_460-532_ alone or the mixture of RabL2 and IFT81_460-533_/74_460-532_. The results show that IFT81_460-533_/74_460-532_ in the absence of RabL2 does not stimulate hydrolysis of GTP but in the presence of RabL2, the GTP hydrolysis rate is increased by approximately 5-fold when compared to RabL2 alone (Fig. 3D). While highly significant, this 5-fold increase in GTPase activity is less than what was observed with longer IFT81/74 constructs in context of the IFT-B1 pentamer (Fig. 2D). A likely explanation for the lower activity could be that parts of the IFT81/74 complex other than the 70-residue coiled-coil region are required for the optimal positioning of residues involved in GTP hydrolysis. Alternatively, it could be that IFT81_460-533_/74_460-532_ in isolation does not adopt a perfectly productive conformation to allow for the full stimulation of GTPase activity. In any case, IFT81_460-533_/74_460-532_ increases the GTPase activity of RabL2 and likely constitutes the main high-affinity binding site for RabL2 within the IFT-B complex.

### Structural modelling of a minimal IFT81/74/RabL2 complex

To obtain structural insights into the binding of RabL2 to the IFT81/74 complex, we carried out structural modelling using Alphafold (Jumper *et al*., 2021). Previously determined crystal structures are published for the IFT27/25 complex (Bhogaraju *et al*., 2011) and the N-terminal parts of IFT81/74 in complex with GTP-bound IFT22 (Wachter *et al*., 2019). However, no structural information is available for the C-terminal parts of IFT81/74 that we mapped as the binding site for RabL2. Modelling was carried out using the original Alphafold2 algorithm (Jumper *et al*., 2021) as well as the later published Alphafold multimer (Evans *et al*., 2022) using the IFT81_460--533_/74_460-532_ truncation and full length RabL2 for both human and *Chlamydomonas* complexes (Fig. 4A-B). The structural models of the RabL2-bound IFT81_460-533_/74_460-532_ were consistently predicted with high confidence and low error in relative positioning of subunits within the complex (Fig. S4). The core GTPase domain of RabL2 is predicted with very high confidence although the very N- and C-termini of RabL2 are predicted with low confidence and likely displays a high degree of structural flexibility (Fig. S4). Models of the human and *Chlamydomonas* RabL2-IFT81_460-533_/74_460-532_ complexes reveal a similar architecture with a highly similar binding mode for RabL2 on IFT81/74 (Fig. 4A-B). In the structural model shown in Fig. 4C, the RabL2 switch regions are highlighted and nonhydrolyzable GTP analogue GppNHP and Mg^2+^ are modelled based on the crystal structure of Rab8 (PDB code 4LHW, cite (Guo *et al*., 2013)). Interestingly, although GTP was not part of the structural modelling, the switch regions of RabL2 adopt a conformation very similar to that of other Rab GTPases bound to GTP or GTP analogues. The structural prediction shows that the IFT81/74-RabL2 complex is mainly formed through interactions with the switch regions of RabL2 and IFT74 with fewer contacts to IFT81 (Fig. 4C). We note that this binding mode is consistent with IFT81/74 being an effector of RabL2. In the structural model, IFT81/74 does not insert any residues into the active site of RabL2 and no amino acid sidechain of IFT81/74 is closer than 10Å from the GTPase site of RabL2. This indicates that the GAP activity of IFT81/74 towards RabL2 does not utilize the insertion of one or more residues *in trans* into the active site of RabL2.

**Figure 4:**
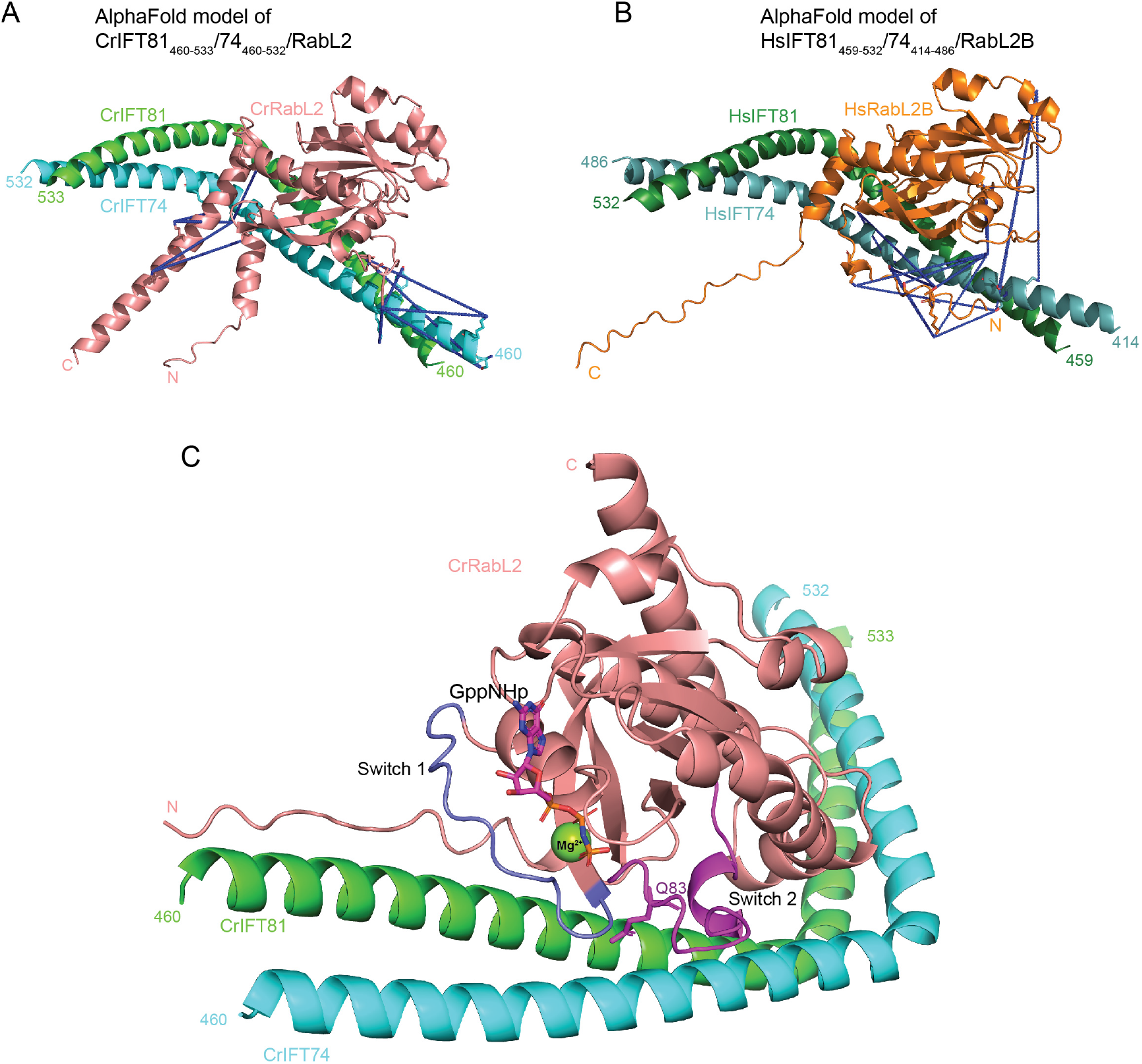
AlphaFold predicted structures of minimal IFT81/74/RabL2 complexes. **(A)** AlphaFold predicted model of a *Chlamydomonas* IFT81_460-533_/74_460-532_/RabL2 complex. The MS identified crosslinks between RabL2, and IFT81/74 are shown as blue dashed lines going between contributing reactive side chains displayed as sticks. **(B)** AlphaFold predicted model of human IFT81_459-532_/74_414-486_/RabL2B with labelled intermolecular crosslinks as in (A). **(C)** Structural model of CrIFT81_459-532_/74_414-486_/RabL2B highlighting the binding of the RabL2 switch 1 (colored slate) and 2 (colored magenta) regions with IFT81/74. The nonhydrolyzable GTP analog GppNHp is shown as sticks and the Mg^2+^ ion as a sphere after superpositioning the Rab8-structure (pdb code 4LHW) onto CrRabL2.

The binding site for RabL2 on IFT81/74 was further confirmed by chemical cross-linking coupled to mass spectrometry (XL-MS). Both human and *Chlamydomonas* IFT-B1 hexamers were chemically crosslinked using the amine- and hydroxy-specific, homo-bifunctional and MS-cleavable crosslinker disuccinimidyl dibutyric urea (DSBU) with a crosslinking space arm of 12.5Å (Iacobucci *et al*., 2018). MS-cleavable crosslinkers can be cleaved in the mass spectrometer yielding two linear peptides which can be then easily identified. Protein complexes were crosslinked by incubation with 0.25mM DSBU and then digested with both LysC and trypsin. After digestion and to increase their identification rates, cross-linked peptides were enriched by strong cation-exchange chromatography (SCX) and then subjected to MS/MS analysis. Using MeroX software (Götze *et al*., 2015) we identified 137 intra- and 211 intermolecular cross-links at an FDR of 1% for the *Chlamydomonas* IFT-B1 hexamer of which 34 belong to RabL2. For the human IFT-B1 hexamer, 229 intra- and 333 intermolecular cross-links were identified, of which 25 belong to RabL2. For a more comprehensive analysis, the intermolecular crosslinking pairs formed between RabL2 and IFT81 or IFT74 were mapped on the AlphaFold predicted structures as solid lines (Fig. 4A-B, Movies 1-2 and Sup. Table 1). The MS/MS cross-linking data are consistent for human and *Chlamydomonas* complexes and show that reactive residues in RabL2 mainly cross-link to residues of IFT74 with fewer cross-links to IFT81 (Movies 1-2). Most of the reactive side chains of the RabL2-IFT74 and RabL2-IFT81 crosslinking pairs are located close to each other within a distance of 25Å with only 3 crosslinking pairs formed 36-43Å apart. Human RabL2B crosslinks along a 49Å long surface formed by coil-coils of IFT74_428-458_ and IFT81_467-500_ whereas the *Chlamydomonas* counterpart crosslinks only to the IFT74_460-506_ helix comprising a 64Å long surface. The chemical crosslinking results provide a validation of the computational predictions of the IFT81/74-RabL2 complex structures and corroborate a conserved binding site for RabL2 on IFT81/74.

### *Chlamydomonas* IFT81/74 binds human RabL2 to stimulate GTP hydrolysis

All subunits of the core IFT machinery, including IFT81, IFT74 and RabL2 are conserved between *Chlamydomonas* and human (van Dam *et al*., 2013), which is quite astonishing given the more than 1B years of evolution separating the two species (Dutcher *et al*., 2012). CrRabL2 and HsRabL2 share 49% identity on the amino acid level. Structural models of *Chlamydomonas* and human RabL2-IFT81_460-533_/74_460-532_ shown in Fig. 4A-B suggest a common binding site for RabL2 on the IFT81/74 complex. To assess the conservation of the RabL2 binding site on IFT81/74, the ConSurf server (Glaser *et al*., 2003; Landau *et al*., 2005) was used to plot the amino acid conservation onto the surface of the structural model (Fig. 5A-B). The conservation plot reveals that the residues of IFT81/74 involved in binding RabL2 are highly conserved across ciliated species containing RabL2 (Fig. 5B). Interestingly, in *C. elegans*, where RabL2 is absent, the RabL2 binding site on IFT81/74 is poorly conserved and partially missing (Fig. S5). The surface of RabL2 engaging in interactions with IFT81/74 is also well conserved albeit less so than the IFT81/74 surface (Fig. 5B).

**Figure 5:**
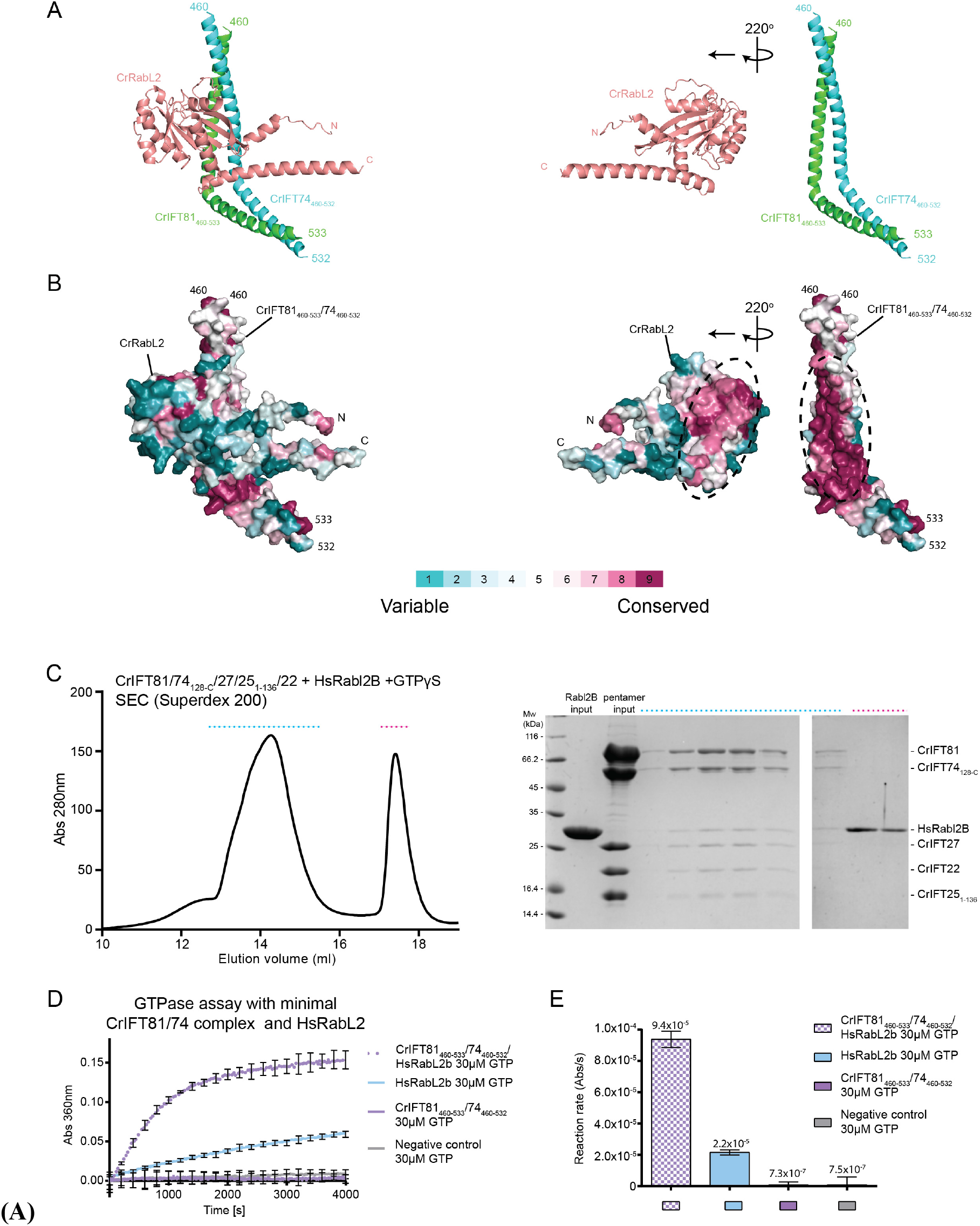
RabL2 interacts with IFT81/74 through a conserved interface. **(A)** The predicted Cr IFT81_460-533_/74_460-532_/RabL2 structure (left) is computationally ‘opened up’ by rotating RabL2 220° around the y-axis (right) to visualize the binding interface between RabL2 and IFT81/74. **(B)** Surface conservation map of the IFT81/74/RabL2 complex in the same position as in (A). Amino acid conservation is indicated according to the color code. **(C)** SEC profile showing that human RabL2B co-purifies with *Chlamydomonas* IFT-B1 pentamer (left). Coomassie stained SDS gel of SEC fractions highlighted with dashed lines on the SEC profile. **(D)** GTPase assay showing that the minimal IFT81/74 complex from *Chlamydomonas* induces GTP hydrolysis in human RabL2B. The curves are averages of 3 independent experiments and error bars shown the standard deviation for each 200s. **(E)** quantification of the reaction rates of the GTPase reaction based on linear regression for the first 500 s of the experiment showing standard deviation as error bars.

To address the highly conserved IFT81/74-RabL2 interface biochemically, we incubated the IFT-B1 pentamer from *Chlamydomonas* with human RabL2 in the presence of GTPγS and carried out SEC. The result shows that human RabL2 indeed co-purifies with the *Chlamydomonas* IFT-B1 pentamer demonstrating a direct physical interaction (Fig. 5C). Co-migration of subunits as a complex on the Superdex 200 column, in our experience, indicates a relatively strong interaction with a Kd in the single digit μM range or lower (Vetter *et al*., 2015b). The high degree of conservation of surface areas of the structural model shown in Fig. 5B thus translates into a biochemically conserved interaction between RabL2 and IFT81/74 across species (Fig. 5C). These results raise the question of whether the mechanism of RabL2 GTP hydrolysis activation by IFT81/74 is also conserved from *Chlamydomonas* to human. To test this, the *Chlamydomonas* IFT81_460-533_/74_460-532_ complex, constituting the minimal binding site for RabL2, was incubated with human RabL2B and GTP in a GTPase assay. The results show that *Chlamydomonas* IFT81_460-533_/74_460-532_ increases the reaction rate of GTP hydrolysis more than 4-fold compared to human RabL2B alone (Fig. 5D). Interestingly, the intrinsic hydrolysis rate of human RabL2B is 29 times higher than that of GTP in buffer and approximately 3 times higher than the intrinsic GTP hydrolysis rate of CrRabL2 (compare Figs. 5D and 3E). However, the >4-fold stimulation of GTPase activity of HsRabL2 by *Chlamydomonas* IFT81_460-533_/74_460-532_ is not too different from the 5-fold increase in activity for CrRabL2. This shows that the RabL2 binding site and the ability to stimulate GTP hydrolysis of RabL2 are conserved between IFT81/74 from *Chlamydomonas* and human. The GAP activity of the IFT-B1 complex, which inactivates RabL2 and dissociates it from the IFT trains, is thus likely to be an ancient mechanism involved in IFT initiation that is conserved across ciliated RabL2-containing organisms.

## Discussion

Here, we map the binding sites for RabL2 on CEP19 and the IFT-B1 complex. We show that the IFT-B1 complex stimulates GTP hydrolysis in RabL2 and map the functional binding site to a short coiled-coil segment of the IFT81/74 hetero-dimer. The intrinsic GTPase activity of RabL2 is increased by about 20-fold when incorporated into the IFT-B1 complex. The intrinsic GTPase activity of small GTPases is typically very low and not compatible with biological timeframes of the processes that they regulate (Cherfils and Zeghouf, 2013), which is also the case for RabL2 (Figs. 1E and 2F-G). GAPs are thus required to stimulate GTP hydrolysis and inactivate the GTPase at the appropriate time and location in the cell. The typical rate enhancement by GAPs are 3-5 orders of magnitude *in vitro*, ensuring almost instantaneous GTP hydrolysis upon complex formation between GTPase and its GAP (Scheffzek and Ahmadian, 2005). The increase in the GTP hydrolysis rate can, however, be a little as 10 times as seen for Sar1 during the dynamic assembly and disassembly of COPII (Antonny and Schekman, 2001). Sar1-GTP initiates coat assembly by associating with vesicles budding from the ER, triggering the subsequent recruitment of Sec23/24 followed by Sec13/31 to complete COPII coat formation. The assembled COPII coat is a GAP for Sar1 that, mainly through Sec23 but assisted by Sec31, activates GTP hydrolysis by Sar1 thus initiating coat disassembly (Bi, Corpina and Goldberg, 2002; Bi, Mancias and Goldberg, 2007). The relatively low activation rate of Sec23/31 towards Sar1 of only one order of magnitude likely reflects the timescale of COPII coat assembly/disassembly, which is on the order of seconds (Antonny and Schekman, 2001). Given that IFT train assembly at the ciliary base likely takes 3-10s before injection into the cilium (Wingfield *et al*., 2017), the relatively low 20-fold activation of the reaction rate for GTP hydrolysis in RabL2 by the IFT complex appears to be biologically meaningful. We note that a much higher GAP activity of the IFT complex towards RabL2 would result in premature dissociation of RabL2 from IFT trains, which would be unproductive. On the other hand, a much lower GAP activity would likely result in too slow GTP hydrolysis in RabL2 and would result in continued association of RabL2 with IFT trains and faulty retrograde transport of BBSomes and associated cargoes as exemplified by the HsRabL2 Q80L mutant defective in GTP hydrolysis (Duan *et al*., 2021).

The wide range of catalytic activation mechanisms of GTP hydrolysis of small GTPases is mirrored by a high degree of structural and functional diversity among different GAPs. Many GAPs do, however, function by inserting one or more residues into the active site of the small GTPase to promote catalysis (Scheffzek and Ahmadian, 2005; Mishra and Lambright, 2016). The archetypical RasGAP functions by inserting an arginine finger into the active site of Ras to neutralize the build-up of negative charge of the transition state (Pan *et al*., 2006; Scheffzek and Shivalingaiah, 2019). Other GTPase families such as Rho and Arl/Arf families also rely on an arginine-finger supplied *in trans*. Interestingly, Rap1GAP works by inserting a catalytic asparagine thumb into the active site of Rap1 (Daumke *et al*., 2004). Rab proteins often rely on GAPs of the TBC domain-containing family where both an arginine finger and a glutamine, replacing the catalytic glutamine of switch 2, are inserted into the active site of the Rab protein (Pan *et al*., 2006). For this reason, mutation of the switch 2 glutamine in some Rabs is not sufficient to create a constitutively active Rab as exemplified by Rab33, which associate with the dual-finger RabGAP RUTBC1 (Nottingham *et al*., 2011; Cherfils and Zeghouf, 2013). In the structural models of the IFT81/74-RabL2 complexes presented in Fig. 4, the IFT81/74 complex mainly associates with the switch regions of RabL2 but do not appear to insert any residues into the GTP-binding active site. There are several examples of GAPs that activate GTP hydrolysis of small GTPases without inserting residues directly into the active site. MnmE and dynamin family GTPases were shown to use K^+^/Na^+^ cations instead of a catalytic arginine (Mishra and Lambright, 2016). In addition, the structure of Ran bound to RanGAP and RanBP1 shows that the Ran protein itself provides the catalytic machinery without the insertion of residues from RanGAP into the active site (Seewald *et al*., 2002). Instead, RanGAP and RanBP1 appear to activate Ran via an allosteric effect that stabilizes the switch regions including switch 2, which contain the catalytic glutamine. Given the structural model in Fig. 4, it appears likely that the IFT-B1 complex stimulates the GTP hydrolysis activity of RabL2 through an allosteric effect that stabilizes a catalytically competent conformation of the switch regions and active site of RabL2. We show that the mechanism of GTP hydrolysis in RabL2 does rely on a classical catalytic switch 2 glutamine as the Q83L mutation abolishes the enhanced GTPase activity of a RabL2_Q83L_ containing IFT-B1 hexamer (Fig. 2F-G). The detailed unravelling of the catalytic mechanism of RabL2 and GAP function of the IFT-B1 complex awaits detailed experimental and structural elucidation.

The data presented here combined with previously published results allow us to propose a model for RabL2 function in IFT initiation (Fig. 6A). In this model, RabL2 is recruited, most likely in the GTP-bound form, to the base of the cilium via an interaction with the centriolar protein CEP19. RabL2-GTP is then handed over to and incorporated into the IFT trains through an interaction with the C-terminal part of the IFT81/74 sub-complex. Interestingly, the effect of CEP19 knockout on ciliogenesis can in part be rescued by over-expressing WT RabL2 and fully rescued by over-expressing RabL2 Q80L (Kanie *et al*., 2017). This result could suggest that the main role of CEP19 in IFT initiation is to concentrate the RabL2 protein at the base of the cilium. In agreement with this, CEP19 knockout in mice is not embryonic lethal but results in morbidly obese and hyperphagic mice (Shalata *et al*., 2013). Interestingly, a homoallelic nonsense mutation in CEP19 gene that leads to premature truncation at residue R82 manifested by obesity, decrease sperm count, fatty lever, heart problems and intellectual disability has been documented in humans (Shalata *et al*., 2013). We note that the human CEP19 protein sequence used by Shalata et al., 2013 has four additional N-terminal residues (MYMG) as compared to the CEP19 sequence available in Unitprot (https://www.uniprot.org/uniprot/Q96LK0). Our data suggest that this CEP19_R82*_ mutant lacks the entire RabL2 binding site (Fig. 1) and is thus unable to recruit RabL2 to the ciliary base, which provide a molecular mechanism for the disease phenotype. Following recruitment of RabL2 by CEP19 at the ciliary base, RabL2-GTP is incorporated into IFT trains, which may prime these trains for initiation of anterograde transport, possibly through a conformational change of the IFT-B1 complex (Fig. 6B). Shortly after departure of IFT trains from the ciliary base (Fig. 6C), the GAP activity of IFT-B1 towards RabL2 induces GTP hydrolysis and dissociation of RabL2 from the anterograde train, which may prime the IFT train for proper retrograde transport, possibly through a reversal of the conformational change induced by RabL2 in the IFT-B1 complex (Fig. 6D). The model shown in Fig. 6 agrees with the observation that the RabL2 S35N mutant unable to bind GTP does not rescue ciliogenesis of the RabL2 knockout (Kanie *et al*., 2017), which shows that the incorporation of RabL2-GTP is required for proper cilium formation. On the other hand, the HsRabL2_Q80L_ mutant, where the IFT-B1 complex is unable to promote GTP hydrolysis in RabL2, does rescue ciliogenesis of RabL2 knockouts (Kanie *et al*., 2017). However, the HsRabL2_Q80L_ mutant accumulates in cilia together with the BBSome protein BBS4 and the GPCR GPR161, which is not the case in RabL2 WT or S35N mutant RabL2 (Kanie *et al*., 2017). This ciliary accumulation of BBSomes and GPCRs let Duan et al., to suggest that RabL2 regulates BBSome mediated ciliary export (Duan *et al*., 2021). In our model presented in Fig. 6, the RabL2_Q80L_ mutant prevents the switch to retrograde IFT by staying associated with IFT trains, which could possibly prevent the association of the BBSome with IFT25/27 during retrograde transport (Eguether *et al*., 2014). We note that RabL2 is located close to IFT27/25 in the C-terminal region of the IFT81/74 sub-complex. The observation that HsRabL2_Q80L_ accumulates BBSome components and GPCRs in cilia may thus be an effect of continued association of RabL2-GTP with IFT trains rather than a direct cellular function of RabL2 in fine-tuning ciliary export of signalling factors.

**Figure 6:**
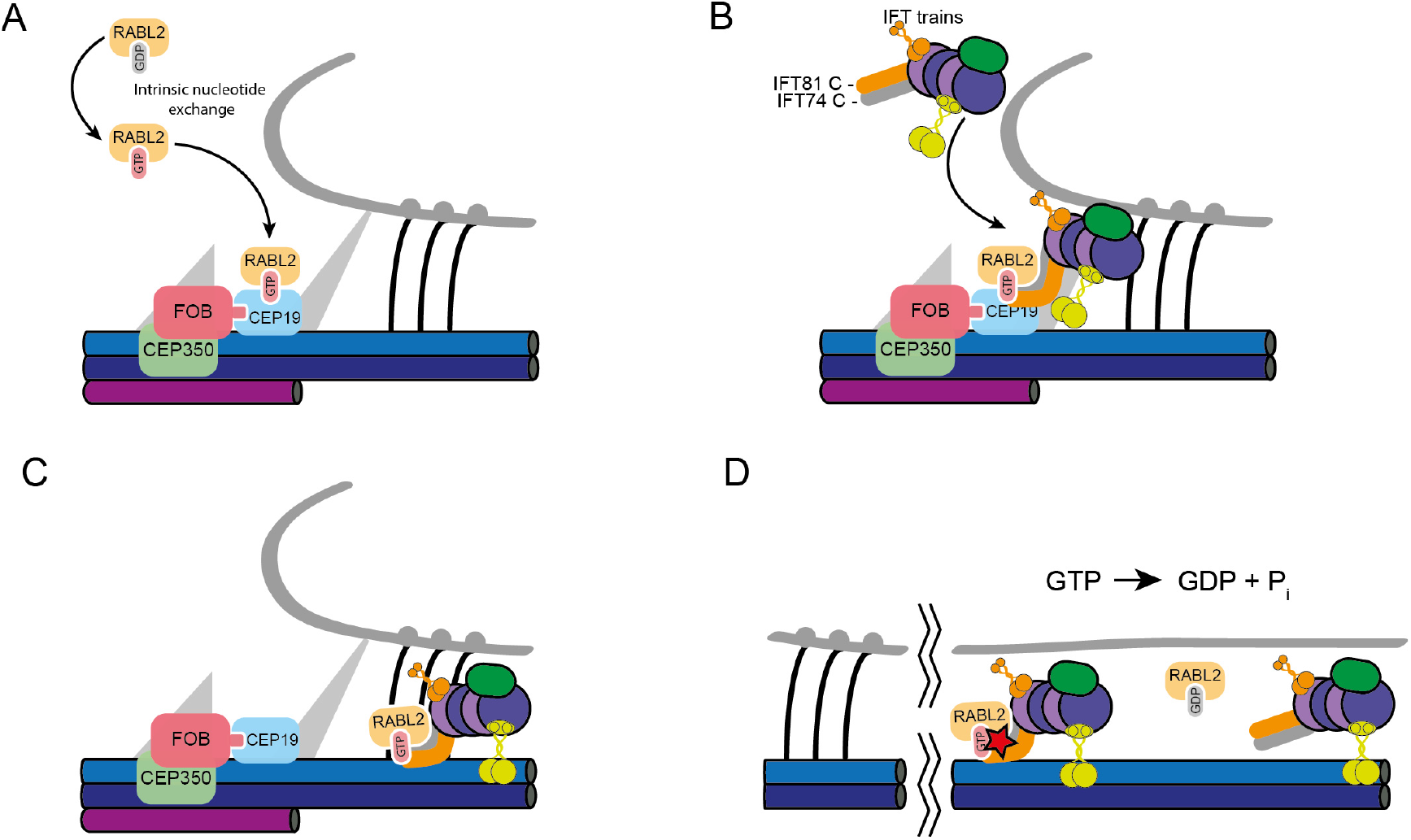
Model for RabL2 function in the initiation of IFT. **(A)** Following activation of GDP-RabL2 by intrinsic nucleotide exchange, active GTP-bound RabL2 is recruited to the ciliary base by CEP19. **(B)** GTP-RabL2 is then handed over to the IFT train by interaction with a stretch of 70 residues of the coiled-coil domain of IFT81/74. **(C)** The association between GTP-RabL2 and the IFT81/74 initiates ciliary entry of anterograde IFT trains together with GTP-RabL2. **(D)** Once in the cilium, GTP-hydrolysis in RabL2 is stimulated by IFT81/74 converting active GTP-bound RabL2 into inactive GDP-bound RabL2. Inactive GDP-RabL2 then dissociates from the anterograde IFT train.

## Materials and Methods

### Cloning and expression of CEP19 and RabL2 in *E.coli*

The gene encoding *C. reinhardtii* CEP19_1-208_ was subcloned from the pGEX-6P-1-CrCEP19 to the pEL-K vector. The genes encoding CrRabL2 and HsRabL2B were subcloned from the pGEX-6P-1-CrRabL2 and pGEX-6P-1-HsRabL2B to the pEL-A vector. Protein encoding genes were amplified using forward primer encoding a 5’-His_(6)_-TEV overhang allowing cloning of a N-terminal His_(6)_-TEV-tag and revers primer to allow cloning into the pEL-K vector. The DNA fragments pEL-K, the amplified insert, and the His_(6)_-TEV-tag were assembled by Gibson Assembly (Gibson *et al*., 2009). CrRabL2_Q83L_ mutant was constructed by PCR mutagenesis using pEL-A-His_(6)_-TEV-CrRabL2 as template. The CrRabL2_Q83L-Δ2-8_ truncation was constructed by mutagenesis using pEL-A-His_(6)_-TEV-CrRabL2_Q83L_ as template. The primers were constructed to omit 24bp of CrRabL2_Q83L_ starting from the second codon. The protein expressed from this plasmid is hereafter termed CrRabL2_Q83L-9-C_. CrCEP19_107-195_ truncation was constructed by PCR mutagenesis using pEL-K-His_(6)_-TEV-CrCEP19_1-208_ as template. The forward primer omitted the first 318bp of the gene encoding CrCEP19_1-208_ and the revers primer omitted the last 39bp. The forward primer encoding a 5’-His_(6)_-TEV overhang allowing cloning of a N-terminal His_(6)_-TEV-tag and revers primer to allow cloning into the pEL-K vector. Plasmids were transformed into *E.coli* BL21 (DE3) cells by heat shock at 42°C for 40sec. Heat shocked cells were plated on LB agar plates supplemented with appropriate antibiotics and incubated overnight.

For recombinantly expression of CrRabL2, CrRabL2_Q83L_, CrRabL2_Q83L-9-C_, HsRabL2B, CrCEP19_1-208_, CrCEP19_107-195_ an overnight preculture was used to inoculate 6L Terrific broth (TB) medium supplemented with antibiotics. Expression cultures were grown at 37°C until OD_600_ reached 1.2, at this point the temperature was lowered to 18°C and 0.5mM of isopropyl β-D-1-thiogalactopyroside (IPTG) was added after 1.5h. Expression cultures were incubated at 18°C for 18h after induction and the cells harvested by centrifugation (rotor F9-6x1000lex) at 7822 RFC (Relative Centrifugal Force) at 4°C for 12min.

### Purification of CEP19 and RabL2 proteins

*E.coli* cells with over-expressed CEP19 or RabL2 protein were resuspended in lysis buffer (50mM Tris pH 7.5, 150mM NaCl, 10% (v/v) glycerol, 5mM MgCl_2_, 20mM imidazole 5mM BME) containing 1mM phenylmethanesulfonyl fluoride (PMSF) and lysed by sonication using the Sonopuls (Bandelin) fitted with VS 70T probe in two cycles of 10min at 40% amplitude and of 1/1s pulses. The cell lysate was centrifuged (rotor A27-8x50) at 69028 RFC in 4◦C for 30min and the cleared lysate was aspirated and added 1µl DNase (ThermoFisher). Further purification of the CrRabL2, CrRabL2_Q83L_, CrRabL2_Q83L-9-C_, HsRabL2B, CrCEP19_1-208_, CrCEP19_107-195_ were performed by loading the cleared lysate onto an IMAC cOmplete His-Tag 5ml column (Roche) pre-equilibrated in 5 column volumes (CV) of lysis buffer. The column was after loading washed with 5CV of wash buffer (50mM Tris pH 7.5, 1M NaCl, 10% (v/v) glycerol, 5mM MgCl_2_, 20mM imidazole, 5 mM BME) followed by equilibration in 5CV of lysis buffer. Elution was performed on an Äkta primer (GE Healthcare) using a gradient from 0-100% Ni-elution buffer (50mM Tris pH 7.5, 150mM NaCl, 10% (v/v) glycerol, 5mM MgCl_2_, 600mM imidazole, 5 mM BME). Peak fractions were pooled and 1mg of TEV (*Tobacco Etch Virus*) protease was added followed by overnight dialysis at 4°C against dialysis buffer (20mM Tris pH 7.5, 50mM NaCl, 10% (v/v) glycerol, 5mM MgCl_2_, 1mM DTT) to cleave the affinity tags. Dialysis sample was loaded onto the cOmplete His-Tag column pre-equilibrated in 5CV of low salt buffer (20mM Tris pH 7.5, 50mM NaCl, 10% (v/v) glycerol, 5mM MgCl_2_, 5mM BME) and for RabL2 proteins, the flow through loaded onto a HiTrap Q HP 5ml anion exchange column (GE Healthcare) pre-equilibrated in low salt buffer. Elution was performed on the Äkta prime with gradient going from 0-100% high salt buffer (20mM Tris pH 7.5, 1M NaCl, 10% (v/v) glycerol, 5mM MgCl_2_, 5mM BME). For CrCEP19_1-208_, CrCEP19_107-195_ the flow through was loaded onto anion exchange MonoQ column (GE Healtcare) pre-equilibrated in low salt buffer. Elution was performed with gradient going from 0-40% high salt buffer on the Äkta purifier (GE Healtcare). Following purification by anion exchange was the protein sample loaded onto the HiLoad 16/600 Superdex 75 (GE Healtcare) on the Äkta purifier pre-equilibrated in SEC buffer (10mM HEPES pH 7.5, 150mM NaCl, 5mM MgCl_2_, 1 mM DTT). Note all CrCEP19 purification buffers were prepared without MgCl_2_.

### Expression and purification of *Chlamydomonas* IFT-B1 complexes

Recombinant expression of *Chlamydomonas* IFT-B1 complexes were achieved by co-expressing plasmids in *E. coli* BL21 (DE3) cells. The CrIFT81/74_128-C_/27/25_1-136_/22 complex was expressed from pEC-A-His_(6)_-TEV-CrIFT81, pEC-K-His_(6)_-TEV-CrIFT74_128-C_, pEC-S-His_(6)_-TEV-CrIFT25_1-136_, pEC-Cm-CrIFT27 and pEC-A-CrIFT22. The CrIFT81_152-C_/74_150-C_/27/25_1-136_ complex was expressed from pEC-A-His_(6)_-TEV-CrIFT81_152-C_, pEC-K-His_(6)_-TEV-CrIFT74_150-C_, pEC-S-His_(6)_-TEV-CrIFT25_1-136_ and pEC-Cm-CrIFT27. The CrIFT81_133-475_/74_132-475_/22 complex was expressed from pEC-A-His_(6)_-TEV-CrIFT81_133-475_, pEC-K-His_(6)_-TEV-CrIFT74_132-475_ and pEC-A-CrIFT22. The CrIFT81_460-C_/74_460-C_/27/25_1-136_ complex was expressed from pEL-A-His_(6)_-CrIFT81_460-C_, pEL-K-His_(6)_-Strep-TEV-CrIFT74_460-C_ and pEC-S-His_(6)_-TEV-CrIFT25_1-136_-RBS-CrIFT27. The CrIFT81_460-623_/74_460-615_/27/25_1-136_ complex was expressed from pEL-A-His_(6)_-CrIFT81_460-623_, pEL-K-His_(6)_-TEV-CrIFT74_460-615_ and pEC-S-His_(6)_-TEV-CrIFT25_1-136_-RBS-CrIFT27. The CrIFT81_460-533_/74_460-532_ complex was expressed from pEL-A-His_(6)_-CrIFT81_460-533_ and pEL-K-His_(6)_-TEV-CrIFT74_460-532_. For recombinantly expression of IFT-B1 complexes an overnight preculture was used to inoculate 6L Terrific broth (TB) medium supplemented with antibiotics. Expression cultures were grown at 37°C until OD_600_ reached 0.8, at this point the temperature was lowered to 18°C and 0.5mM of IPTG was added after 1.5h. Expression cultures were incubated at 18°C for 18h after induction. Lysing cells expressing CrIFT81/74_128-C_/27/25_1-136_ allowed for the purification of the tetrameric IFT-B1 complex, co-lysing these with cells expressing IFT22 and/or CrRabL2 yields pentameric or hexameric IFT-B1 complexes, respectively. Harvest and sonication of cells were performed as described above for RabL2 and CEP19 species. Cleared lysate was loaded onto the cOmplete His-Tag column pre-equilibrated in lysis buffer (50mM Tris pH 7.5, 150mM NaCl, 10% (v/v) glycerol, 20mM imidazole, 1mM MgCl_2_, 5mM BME), followed by 5CV wash buffer (50mM Tris pH 7.5, 1M NaCl, 10% (v/v) glycerol, 20mM imidazole, 1mM MgCl_2_, 5 mM BME). The column was washed with low salt buffer (50mM Tris pH 7.5, 75mM NaCl, 10% (v/v) glycerol, 1mM MgCl_2_, 5mM BME). Elution was performed directly from the cOmplete His-tag column through the HiTrap Hp Q anion column with Ni-elution buffer (50mM Tris pH 7.5, 75mM NaCl, 10% (v/v) glycerol, 600mM imidazole, 1mM MgCl_2_, 5mM BME). The elution was collected and loaded onto the HiLoad 16/600 Superdex 200 (GE Healthcare) equilibrated in SEC buffer (10mM HEPES pH 7.5, 150mM NaCl, 1mM MgCl_2_, 1mM DTT).

### Cloning, expression, and purification of human IFT81/74/27/25/22 complex from SF21 insect cells

A donor plasmid that contained the genes encoding for the HsIFT81/74/27/22 complex was designed and purchased from VectorBuilder®. The commercially available pFastBac™ Dual plasmid was used as a template. The five genes were constructed on two opposing multiple cloning sites flanked by Tn7L and Tn7R elements. In one cloning site resided the IFT81 gene preceded by the p10 promoter and succeeded by the SV40 poly-adenylation (pA) early terminator sequence followed by the polyhedrin promoter (pH), IFT74 gene and Tk pA terminator. In the other cloning site, it was constructed a succession of DNA sequences as follows: one pH promoter, the IFT27 isoform 1 gene, and the SV40 pA terminator, one pH promoter, IFT25 gene, Tk pA terminator, one pH promoter, IFT22 isoform a, and SV40 early terminator. The final construct, named pNAP-AG-HsIFT81/74/27/25/22, was compatible with the Bac-to-Bac® Baculovirus expression system. The pNAP-AG-HsIFT81/74/27/25/22 plasmid was used for transformation of DH10bac cells and white colonies indicating successful transposition were selected. The bacmid DNA of white colonies was purified and the insertion of the HsIFT81/74/27/22 cassette into the attTn7 docking site was confirmed by PCR and sequencing. From the correct bacmid, a recombinant baculovirus was produced in SF21 cells as described previously in (Taschner *et al*., 2016, p. 172).

To allow purification of the HsIFT81/74/27/22 complex, DNA sequences encoding both a deca-histidine tag followed by a TEV-cleavage site at the N-terminus of IFT81 and a hexa-histidine tag at the N-terminus of IFT74 followed by Strep and TEV cleavage, were included. For production of large amounts of the HsIFT81/74/27/25/22 complex, 1-3L of SF21 cells in Gibco™ Sf-900™ II SFM media with a density of 1.4 x 10^6^ cells/ml was infected with 8ml/L of P2 virus and incubated for 72h at 27°C. The infected cells were harvested by centrifugation, resuspended in one volume of lysis buffer (50mM Tris-HCl, pH 7.5, 150mM NaCl, 1mM MgCl_2_, 10% glycerol, 5mM BME, 1x complete protease inhibitor mixture (Roche), 10μg/ml DNase I, weight/ml) and lysed by 50x strokes in a Dounce tissue homogenizer. The cell debris was cleared by centrifugation at 20000 x g for 30min, 4°C and the supernatant was filtered through a 5µm filter. The supernatant from the previous step was loaded onto a pre-equilibrated TALON HiTrap column with a peristaltic pump by recirculation at a 5ml/min flow rate. 4 washing steps were performed; of which the first with only lysis buffer, the second with lysis buffer supplemented with 40mM Imidazole pH 7.5, the third with high salt buffer (50mM Tris-HCl, pH 7.5, 1M NaCl, 1mM MgCl_2_, 10% glycerol, 5mM BME) and the fourth with low salt buffer (50mM Tris-HCl, pH 7.5, 75mM NaCl, 1mM MgCl_2_, 10% glycerol, 5mM BME). The HsIFT81/74/27/25/22 complex was eluted from the TALON with a lysis buffer supplemented with 300mM Imidazole pH 7.5. Elutions were concentrated with a 100kDa cutoff Amicon Ultra filter and loaded onto Superose 6 Increase with a 500µl injection loop. Samples were collected from each SEC fraction, migrated on SDS-PAGE and visualized by staining the gel with 1% of Coomassie Briliant Blue. The SEC fractions that contained of all proteins within HsIFT81/74/27/22 complex were combined and concentrated. Hexameric IFT-B1 complex was obtained by mixing purified HsIFT81/74/27/22 with HsRabL2 and pefeorming SEC on a Superose6 column.

### Pulldown assays

Samples containing purified Rabl2_Q83L_, IFT81_460-533_/74_460-532_ and nucleotides (where indicated) were incubated for 1h, at 4°C in a 150 µl PD buffer (50mM Tris-HCl, pH 7.5, 150mM NaCl, 1mM MgCl_2_, 5mM TCEP). Each sample was added to 30µl pre-equilibrated Ni^2+^-NTA beads and incubated at 4°C for 1h. The beads were recovered by low-speed centrifugation (1100 x g) for 3min and washed three times to remove the unbound proteins. The first washing step was performed with 500µl of PD buffer and the second and the third washing with PD buffer supplemented with 20mM Imidazole pH 7.5. The histidine-tagged IFT81_460-533_/74_460-532_ was eluted from the beads by incubation with 50µl PD buffer supplemented with 600mM Imidazole pH 7.5 for 10min. The protein composition of each sample was evaluated on SDS-PAGE stained with Coomassie Briliant Blue.

### Isothermal titration calorimetry (ITC)

ITC was performed at 25°C using a VP-ITC MicroCalorimeter (MicroCal, GE Healthcare). CrRabL2 and CrCEP19 were buffered in 10mM Hepes pH 7.5, 150mM NaCl, 5mM MgCl_2_, 1mM TCEP, and 1mM GTP. A volume of 1.426ml of 10µM CrRabL2 was titrated with 100µM CrCEP19 over 29 injections of 10µl with a stirring speed of 312rpm. Duration of each injection was 17.1sec with 200sec of spacing between individual injections. Titrations were performed in triplicates, and for each titration a background curve consisting of titrant titrated into buffer was subtracted. The ITC data were analysed with Origin 7 provided by MicoCal. The CrRabL2-IFT-B interaction was analyzed on a VP-ITC MicroCalorimeter (MicroCal) instrument. A volume of 1.426ml of IFT81_152-C_/74_150-C_/27/25_1-136_ supplemented with 100µM GTPγS was titrated with a solution of 100µM Rabl2_Q83L_ supplemented with 100µM GTPγS over 29 injections of 10µl with a stirring speed of 312rpm. Three measurements were performed at 25°C in a buffer containing 20mM Hepes pH 7.5, 150mM NaCl, 1mM MgCl_2_ and 0.5mM TCEP. The obtained ITC data were analysed with the Origin 7 software provided by MicroCal.

### Protein complex prediction with AlphaFold multimer

For predicting the structure of Rabl2 in complex with IFT81/74 or CEP19, we used a local installation of AlphaFold v2.1.0 (Jumper *et al*., 2021; Evans *et al*., 2022) as well as a modified version on Colab notebook (Mirdita et al., 2021). As input for alphafold, we used the relevant protein sequences or truncations from *Chlamydomonas reinhardtii* or *Homo sapiens* IFT81, IFT74, CEP19 and RabL2. All used sequences have >500 homologs in available databases and all structural predictions shown in the figures have low PAE scores for the interacting regions indication a high degree of certainty in the relative positions of subunits within the complexes. All figures of protein structures were prepared using PyMOL v. 2.5 (Schrodinger LLC, https://pymol.org)

### GTPase assays

The GTPase activity was measured using the EnzCheck Phosphate Assay Kit (ThermoFisher) at 22°C. GTPase assays were performed in 200µl volumes in 96-well plates and the release of inorganic phosphate (P_i_) was monitored at 360nm using the MultiSkan Go plate reader (ThermoFisher). Assays were performed in triplicates and a reference data set containing reaction buffer substrate (MESG) and Purine Nucleoside Phosphorylase (PNP) enzyme, was subtracted from each data set. Negative controls contained reaction buffer, MESG, PNP enzyme and GTP, and positive controls contained reaction buffer, MESG, PNP enzyme and 100µM P_i_. Reaction components were mixed and incubated at 22°C for 15min before GTP was added and the 96-well plate was shaken for 2sec in the plate reader before measurement was started. Reaction concentrations of CrRabl2 and CrRabL2/CrCEP19 (Fig. 1E) was 250µM and 1mM GTP,and concentrations of IFT-B1 complexes were 55-70µM and 1mM GTP for Fig. 2E, and 60µM IFT-B1 and 30µM GTP for Figs. 2F-G, 3D and 5D. IFT-B1 complex were incubated at 22°C for 6h followed by buffer exchange to ensure that any co-purified nucleotides were hydrolyzed, and P_i_ was removed before the GTPase was performed.

### DSBU crosslinking and mass spectrometry analysis

Cross-linking experiments were performed using the MS-cleavable cross-linker disuccinimidyl dibutyric urea (DSBU). The optimal concentration of DSBU was determined by titration and SDS-PAGE to allow only the formation of specific crosslinks. Reactions containing 20µg of purified IFT-B1 hexameric complexes dissolved in a buffer containing 20mM Hepes pH 7.5, 150mM NaCl, 1mM MgCl_2_ and 0.5mM tris(2-carboxyethyl)phosphine (TCEP) were crosslinked for 30 minutes with various DSBU concentrations ranging from 0.1mM to 1.53mM (representing a molar excess rage from 15 to 250 times respectively), quenched by addition of 1M Tris pH 8.8 and monitored on SDS PAGE and Coomassie staining. A DSBU concentration of 0.25mM per 6µM of IFT-B1 hexamer was chosen as an optimal crosslinking concentration for the data presented in this work.

### Mass spectrometry analysis of DSBU crosslinked protein complexes

For the *Chlamydomonas* IFT-B1 hexamer, 150μg of the purified complex were cross-linked at 6 μM (1.33mg/mL) and a final DSBU concentration of 0.25mM for 30min at room temperature (RT). The reaction was stopped by adding Tris-HCl pH 8.0 to a final concentration of 125 mM and incubated at RT for additional 10min. The sample was denatured and reduced with 6M guanidine hydrochloride (Guan-HCl) and 10mM TCEP in 200mM Tris-HCl pH 8.3 at 95°C for 5min and stirred at RT for an additional 20min. Cysteine residues were alkylated by incubation with 10mM 2-chloroacetamide (CAA) for 20 minutes in darkness. In-solution digestion was performed by incubation with the endoprotease Lys-C (enzyme:protein ratio 1:100) for 1 hour followed by addition of 20 mM ammonium bicarbonate pH 8.0 to reduce the concentration of Guan-HCl to 2M. The protease trypsin was added at an enzyme:protein ratio of 1:50 and incubated at 37 °C overnight. Digested peptides were acidified by adding 25% trifluoroacetic acid (TFA) to a final concentration of 1%.

For the human IFT-B1 hexamer, 60μg of cross-linked complex were denature, reduced and alkylated in lysis buffer containing 5% SDS, 10mM TCEP and 11mM CAA in 100mM Tris-HCl pH8.5 for 10 minutes at 95^0^C. On-bead protein digestion was performed following the Protein Aggregation Capture (PAC) protocol (Tanveer et al., 2019) with some modifications. Cross-linked proteins were precipitated on 120μg of MagResyn HILIC magnetic particles (Resyn Biosciences, Pretoria, Gauteng, South Africa) in 70% acetonitrile for 20 minutes and washed three times with 95% acetonitrile and two times with 70% ethanol. Magnetic particles were then incubated with Lys-c in 50mM ammonium bicarbonate pH 8.0 at a ratio of 1:200 (with respect to protein) for 1h at RT followed by the addition of trypsin in the same buffer at a ratio of 1:100. Trypsin digestion was carried out over night at 37°C and peptides were recovered by transferring the supernatant to a new tube and acidified with 1% TFA.

Digested and acidified peptides from both human and *Chlamydomonas* complexes were first purified with a Sep-Pak tC18 cartridge (Waters Corporation). Briefly, the cartridge was conditioned by adding 50% and 100% acetonitrile and washed with 0.1% TFA prior to sample loading by gravity. Peptides were washed with 0.1% TFA and eluted using 50% and 80% acetonitrile in 0.1% TFA. Organic solvent was removed using a speedvac and one sixth or one tenth (*Chlamydomonas* or human complexes respectively) of the sample was saved as the total peptide mixture for direct MS analysis. The rest was fractionated by cation exchange to enrich for the multiple-charged cross-linked products as follows: Peptides were diluted in 4% phosphoric acid (H_3_PO_4_) and loaded into an Oasis MCX 1 cc Vac Cartridge (Waters Corporation) previously conditioned by adding methanol and 4% H_3_PO_4_. Peptides were recovered in four different fractions through stepwise elution in 0.5% acetic acid and increasing concentrations of ammonium acetate and methanol (Iacobucci et al., 2018). The flow through was also collected. All fractions were desalted again with a Sep-Pak tC18 cartridge as described above.

Total peptide mixtures as well as fractions were analyzed by an Easy nanoLC system coupled directly to a Thermo Fisher Orbitrap Exploris mass spectrometer. Purified peptides were redissolved in 2% acetonitrile, 0.1% TFA, loaded onto a fused silica capillary column (75μm ID, packed in-house with Reprosil - Pur C18, 1.9μm reserved phase material), equilibrated with solvent A (0.1% formic acid), and separated with a linear gradient of 5-45% solvent B (0.1% formic acid, 95% acetonitrile). Mass spectra were acquired using data-dependent acquisition. Top 15 ions with a charge state between 3 and 8 were selected for HCD fragmentation using stepped normalized collision energy (NCE) of 27, 30 and 33. Cross-linked peptides were identified using the Program MeroX version 2.0 (Gotze et al., 2019). In the Mass Comparison settings, the precursor and fragment ion precisions were set to 5.0 and 10.0 ppm respectively, the S/N ratio to 1.5 and the minimum charge to 3. The RISEUP mode was used, maximum missing ions was set to 1 and the neutral loss of identified fragments was selected. In order to have a highly confident identification of the cross-links, a prescore of 30 was applied, the FDR cut off was set to1% and the cRap database was included.

## Acknowledgements

We would like to thank Miren Itxaso Santiago Vela and Maximilian Stoetzel for contributing to preliminary experiments on RabL2 complexes and Anni Christensen for technical assistance with protein purification. We acknowledge Michael Taschner for initial work on sub-cloning RabL2 constructs and for carefully reading this manuscript. This work was funded by grants from the Novo Nordisk Foundation (grant number NNF15OC00114164) and the Independent Research Fund Denmark (grant no: 1026-00016B) to E.L. N.A.P was supported by a postdoc fellowship from the European Commission (H2020, Grant Agreement number 888322). M.L.L and J.S.A was supported by the Independent Research Fund Denmark (grant no: 8021-00425B) to J.S.A.

---

**Figure S1:**
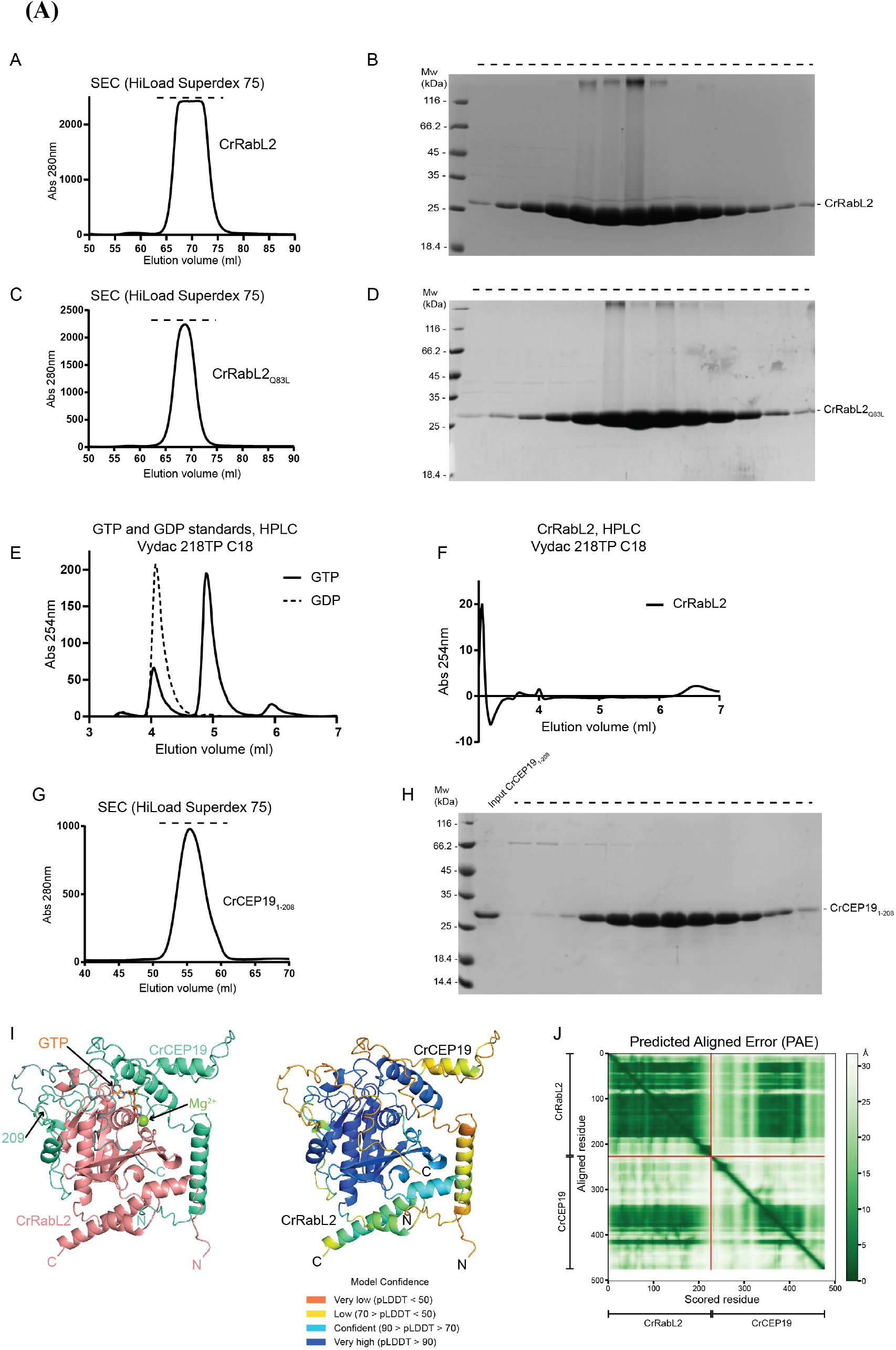
CrRabL2 and CEP19 protein purification and their interaction as predicted by AlphaFold. **(A)** SEC profile of purified CrRabL2. **(B)** The peak fraction highlighted with a top dashed line are verified for purity on SDS PAGE and stained with Coomassie. **(C)** SEC profile of CrRabL2_Q83L_ mutant displaying a similar elution pattern to WT CrRabL2. **(D)** The SEC fractions highlighted with a top dashed line are verified for purity on SDS PAGE and stained with Coomassie. **(E)** High-performance liquid chromatography (HPLC) analysis showing the elution of GTP (solid line) and GDP (dashed line) standards on a Vydac 218TP C18 column. **(F)** HPLC run of purified CrRabL2 demonstrating that the protein does not co-purify with GDP or GTP nucleotides. **(G)** SEC profile of purified CrCEP19_1-208_. **(H)** The SEC CEP19_1-208_ fractions marked by a top dashed line in (G) are migrated on SDS PAGE and stained with Coomassie. **(I)** AlphaFold predicted structural model of CrRabL2 (colored salmon) in complex with CEP19 (green) is shown as cartoon representation on the left. The GTP as well as the position of Mg^2+^ is modelled in the structure after superimposition with the Rab8-GppNHp (PDB code 4LHW). The right panel shows the CrRabL2-CEP19 complex colored according to the pLDDT score. **(J)** Predicted alignment error plot of the CrRabL2-CEP19 complex.

**Figure S2:**
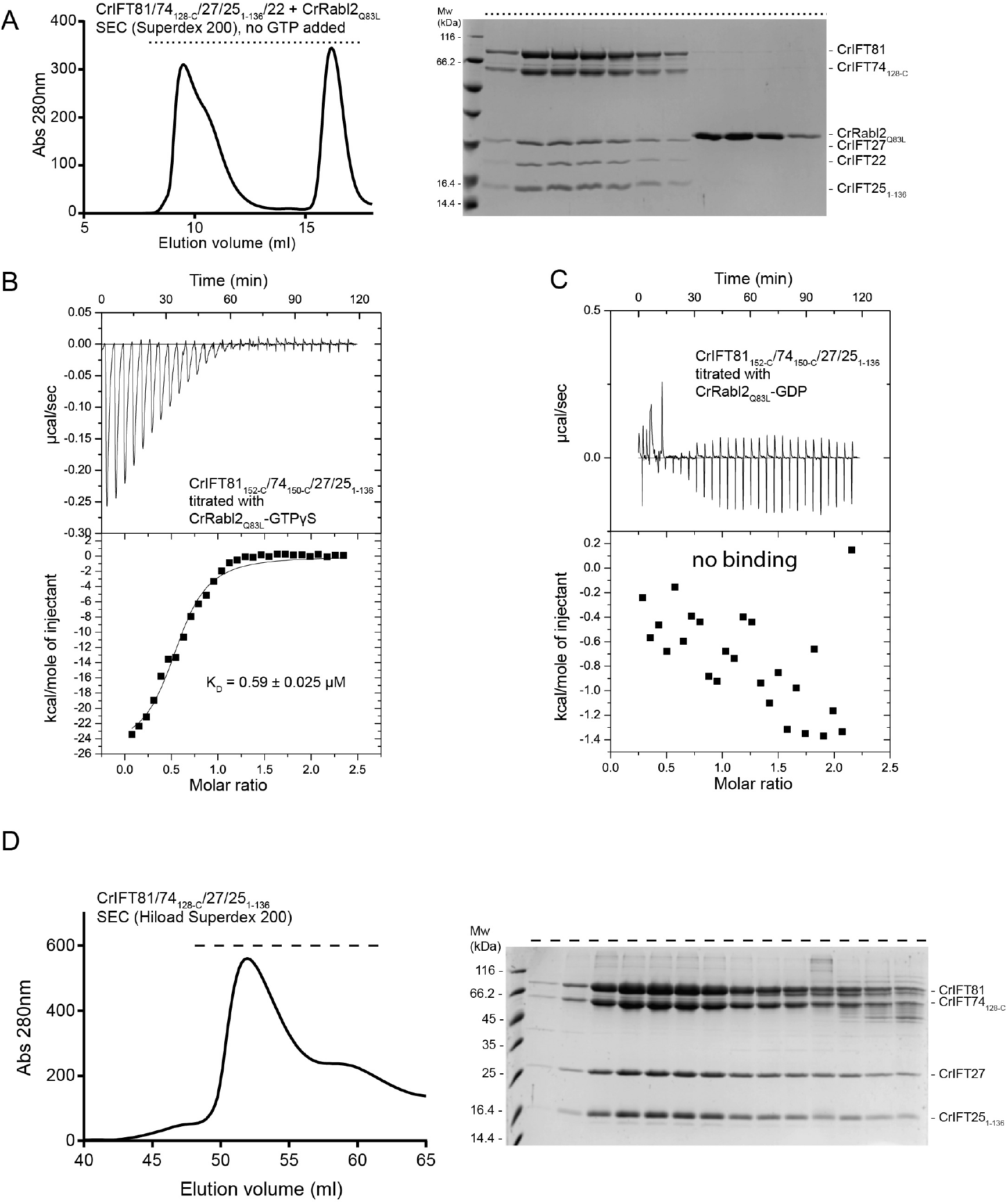
Rabl2 binds to the IFT-B1 tetramer or pentamer only in the presence of GTP. **(A)** Size exclusion chromatogram (left panel) and the corresponding SDS-PAGE gel (right panel) showing that CrRabL2_Q83L_ cannot form a complex with CrIFT81/74_128-C_/27/25_1-136_/22 in the absence of GTPγS. The dashed lines above chromatograms indicate the SEC fractions migrated on SDS-PAGE and stained with Coomassie Brilliant blue. **(B)** ITC measures micromolar affinities of RabL2_Q83L_ for the IFT81_152-C_/74_150-C_/27/25_1-136_ complex in the presence of GTPγS. K_D_ represents the average dissociation constant in µM calculated from three independent experiments. **(C)** No binding is observed for RabL2_Q83L_ to the IFT81_152-C_/74_150-C_/27/25_1-136_ complex in the presence of GDP in ITC. **(C)** SEC profile of IFT-B1 tetramer purification (left) with the corresponding Coomassie staining (right). This sample is used for the GTPase assay shown in fig. 2E-F.

**Figure S3:**
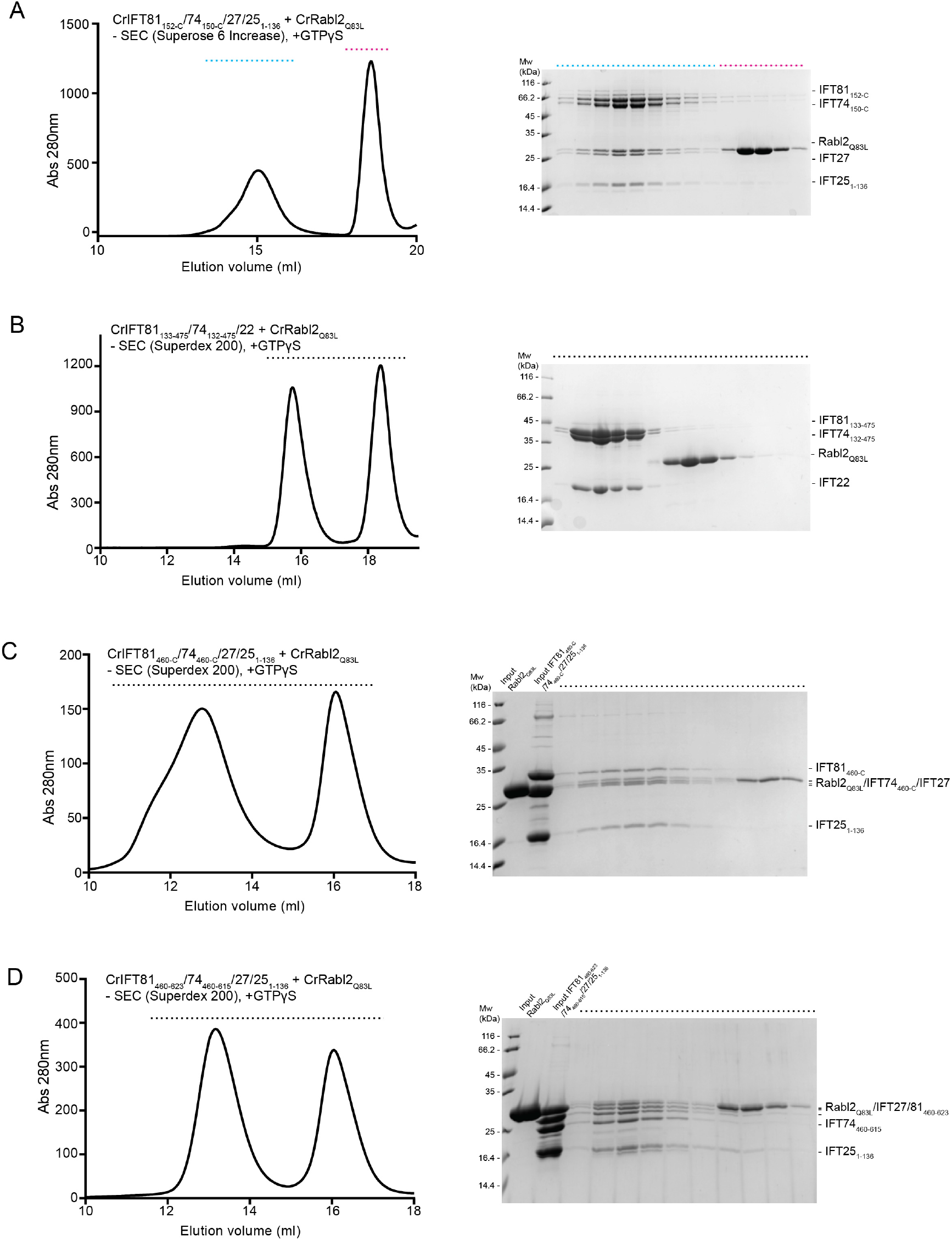
CrRabl2 associates with the C-termini of the CrIFT81/74 complex. (A) A size exclusion chromatogram (left) and the corresponding SDS-PAGE gel (right) that depicts complex formation between CrRabL2_Q83L_ and a tetrameric CrIFT81_152-C_/74_150-C_/27/25_1-136_ complex. The dashed lines indicate the SEC exclusion fractions investigated on SDS-PAGE (right) and stained with Coomassie. **(B)** Size exclusion chromatogram (left panel) and the corresponding SDS-PAGE gels (right panel) showing that CrRabL2_Q83L_ cannot form a complex with CrIFT81_132-475_/74_132-475_/22. The Coomassie staining on the right shows the composition of the highlighted SEC fractions (horizontal top dashed line on the left). **(C)** SEC profile indicating binding of CrRabL2_Q83L_ to a CrIFT81_460-C_/74_460-C_/27/25_1-136_ complex. Samples collected from the highlighted SEC fractions (dashed line) are migrated on SDS PAGE and stained with Coomassie (right panel). **(D)** Binding between CrRabL2_Q83L_ and a CrIFT81_460-623_/74_460-615_/27/25_1-136_ in the presence of GTPγS observed on SEC. The SEC fractions marked with a dashed line are monitored on SDS PAGE and stained with Coomassie on the right panel. All SEC runs shown in this figure were in the presence of 1mM of the non-hydrolysable GTP analogue GTPγS.

**Figure S4:**
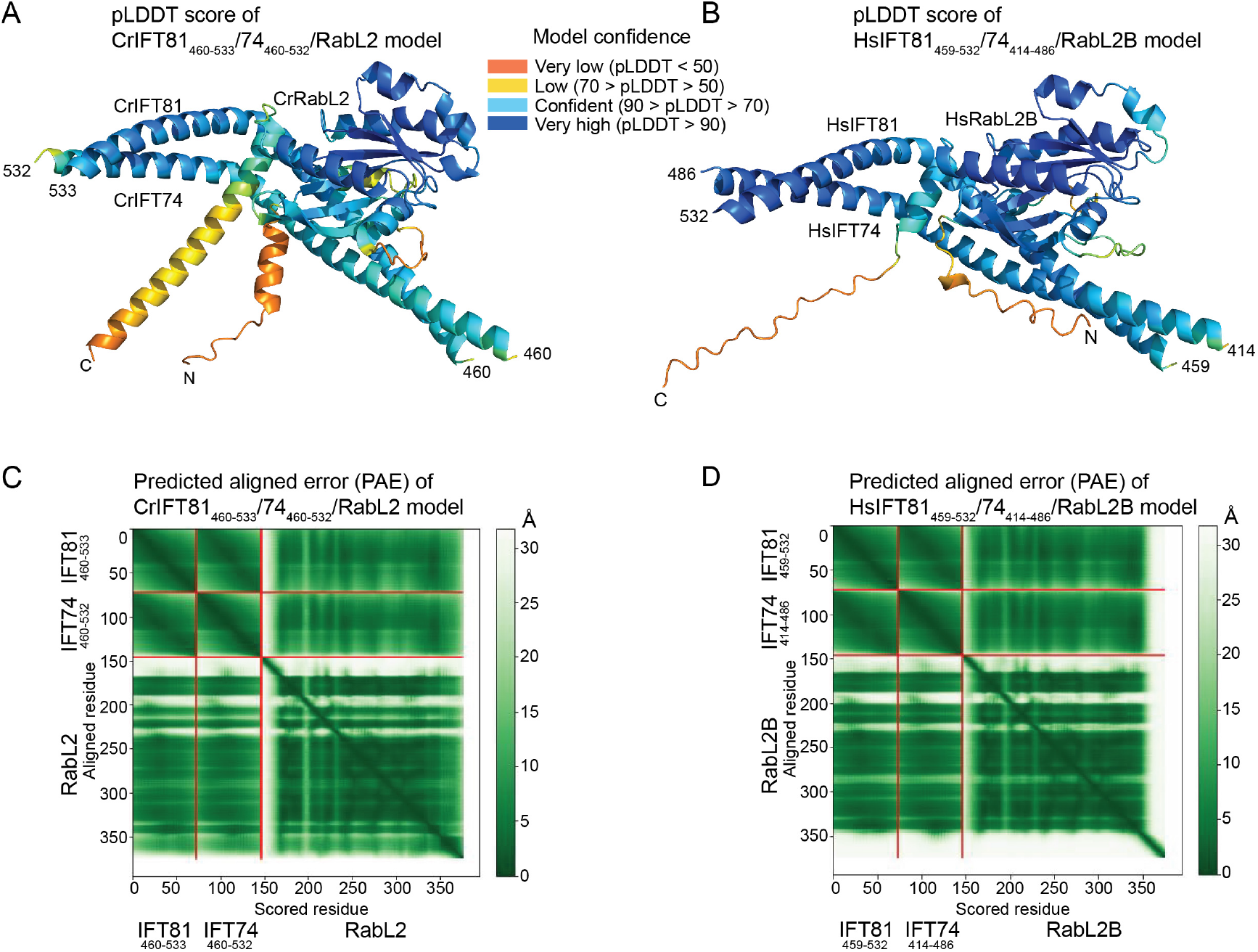
**(A,B)** The AlphaFold models of Cr (A) and Hs (B) minimal IFT81/74/RabL2 complexes in the same orientation as shown in Fig. 4A-B. The structural models are colored by the per-residue estimate pLDDT confidence score. **(C,D)** The plots display the predicted alignment errors (PAE) on a residue basis for the AlphaFold models shown in panels (A) and (B) demonstrating confidence in the relative position of subunits within the complex. The Y-axis show aligned residues and the X-axis show the scored residues. The aligned error is color coded according to the bar to the left of the plots.

**Figure S5:**
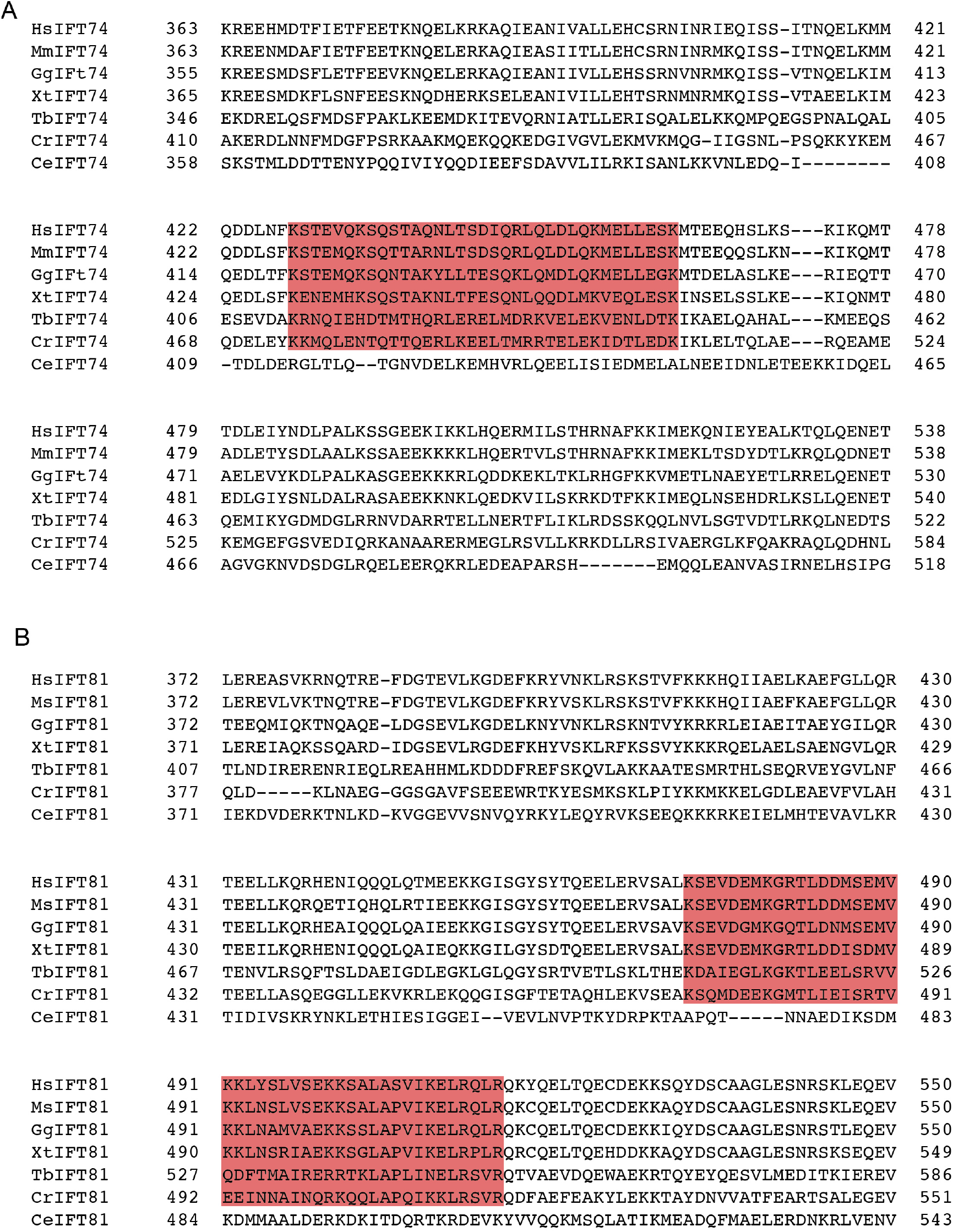
Multiple sequence alignments of IFT74 and IFT81 homologues. Sequence alignment of the C-terminal parts of IFT74 **(A)** and IFT81 **(B)** homologues from Hs (*Homo sapiens*), Mm (*Mus musculus*), Gg (*Gallus gallus*), Xt (*Xenopus tropicalis*), Tb (*Trypanosoma brucei*), Cr (*Chlamydomonas reinhardtii*) and Ce (*Caenorabditis elegans*). RabL2 interacting regions are marked in red boxers.

**Movie 1: 3D representation of the crosslinking network of RabL2 with IFT74_460-C_ and IFT81_460-C_**

The IFT81 is colored in green, IFT74 in cyan and CrRabL2 in salmon. Residues of each protein that contributes to crosslinks are shown as sticks. The intermolecular crosslinks between the reactive side chains are labelled as dashed blue lines.

**Movie 2: 3D representation of the crosslinking network of human RabL2B with human IFT74_414-C_ and IFT81_459-C_**

The HsIFT81 is colored in deep green, IFT74 in teal and human RabL2B in orange. Residues of each protein that contributes to crosslinks are displayed as sticks and the intermolecular crosslinks are labelled as dashed blue lines between the reactive side chains.

**Supplemental material 1: Cross-links identified within the IFT-B1 hexamers by XL-MS/MS**

An excel file with the information obtained with MeroX comprising the crosslinking pairs found within the *Chlamydomonas* and human IFT-B1 hexamers at a high confidence based on a false discovery rate (FDR) of 1%. The “score” is the best score calculated for any combination of the cross-link sites in the two peptides. The “site1” and “site2” represents the crosslinked residues of the first (Prot1) and the second (Prot2) protein. “P%” reflects the probability of the site to be the correct one and “type” indicates whether the crosslinks are taking place within the same protein (intraprotein) or between two different proteins (interprotein). Last, the Crosslinking Spectrum Match (CSMs) represents the number of times that a given crosslinking site has been detected.

## References

Adhiambo, C. et al. (2009) ‘A novel function for the atypical small G protein Rab-like 5 in the assembly of the trypanosome flagellum’, Journal of Cell Science, 122(6), pp. 834–841. doi:10.1242/jcs.040444.

Antonny, B. and Schekman, R. (2001) ‘ER export: public transportation by the COPII coach’, Current Opinion in Cell Biology, 13(4), pp. 438–443. doi:10.1016/S0955-0674(00)00234-9.

Avasthi, P. and Marshall, W. (2013) ‘Ciliary Secretion: Switching the Cellular Antenna to “Transmit”’, Current Biology, 23(11), pp. R471–R473. doi:10.1016/j.cub.2013.04.056.

Bhogaraju, S. et al. (2011) ‘Crystal structure of the intraflagellar transport complex 25/27’, The EMBO Journal, 30(10), pp. 1907–1918. doi:10.1038/emboj.2011.110.

Bhogaraju, S., Engel, B.D. and Lorentzen, E. (2013) ‘Intraflagellar transport complex structure and cargo interactions’, Cilia, 2(1), p. 10. doi:10.1186/2046-2530-2-10.

Bi, X., Corpina, R.A. and Goldberg, J. (2002) ‘Structure of the Sec23/24–Sar1 pre-budding complex of the COPII vesicle coat’, Nature, 419(6904), pp. 271–277. doi:10.1038/nature01040.

Bi, X., Mancias, J.D. and Goldberg, J. (2007) ‘Insights into COPII Coat Nucleation from the Structure of Sec23•Sar1 Complexed with the Active Fragment of Sec31’, Developmental Cell, 13(5), pp. 635–645. doi:10.1016/j.devcel.2007.10.006.

Buisson, J. et al. (2013) ‘Intraflagellar transport proteins cycle between the flagellum and its base’, Journal of Cell Science, 126(1), pp. 327–338. doi:10.1242/jcs.117069.

Cherfils, J. and Zeghouf, M. (2013) ‘Regulation of Small GTPases by GEFs, GAPs, and GDIs’, Physiological Reviews, 93(1), pp. 269–309. doi:10.1152/physrev.00003.2012.

Chien, A. et al. (2017) ‘Dynamics of the IFT machinery at the ciliary tip’, eLife. Edited by A. Akhmanova, 6, p. e28606. doi:10.7554/eLife.28606.

Cole, D.G. et al. (1998) ‘Chlamydomonas Kinesin-II–dependent Intraflagellar Transport (IFT): IFT Particles Contain Proteins Required for Ciliary Assembly in Caenorhabditis elegans Sensory Neurons’, Journal of Cell Biology, 141(4), pp. 993–1008. doi:10.1083/jcb.141.4.993.

van Dam, T.J.P. et al. (2013) ‘Evolution of modular intraflagellar transport from a coatomer-like progenitor’, Proceedings of the National Academy of Sciences, 110(17), pp. 6943–6948. doi:10.1073/pnas.1221011110.

Dateyama, I. et al. (2019) ‘RABL2 positively controls localization of GPCRs in mammalian primary cilia’, Journal of Cell Science, 132(2), p. jcs224428. doi:10.1242/jcs.224428.

Daumke, O. et al. (2004) ‘The GTPase-activating protein Rap1GAP uses a catalytic asparagine’, Nature, 429(6988), pp. 197–201. doi:10.1038/nature02505.

Deane, J.A. et al. (2001) ‘Localization of intraflagellar transport protein IFT52 identifies basal body transitional fibers as the docking site for IFT particles’, Current Biology, 11(20), pp. 1586–1590. doi:10.1016/S0960-9822(01)00484-5.

Ding, X. et al. (2020) ‘Variants in RABL2A causing male infertility and ciliopathy’, Human Molecular Genetics, 29(20), pp. 3402–3411. doi:10.1093/hmg/ddaa230.

Dong, B. et al. (2017) ‘Chlamydomonas IFT25 is dispensable for flagellar assembly but required to export the BBSome from flagella’, Biology Open, 6(11), pp. 1680–1691. doi:10.1242/bio.026278.

Duan, S. et al. (2021) ‘Rabl2 GTP hydrolysis licenses BBSome-mediated export to fine-tune ciliary signaling’, The EMBO Journal, 40(2), p. e105499. doi:10.15252/embj.2020105499.

Dutcher, S.K. et al. (2012) ‘Whole-Genome Sequencing to Identify Mutants and Polymorphisms in Chlamydomonas reinhardtii’, G3 Genes|Genomes|Genetics, 2(1), pp. 15–22. doi:10.1534/g3.111.000919.

Dutcher, S.K. (2014) ‘The awesome power of dikaryons for studying flagella and basal bodies in Chlamydomonas reinhardtii’, Cytoskeleton, 71(2), pp. 79–94. doi:10.1002/cm.21157.

Eguether, T. et al. (2014) ‘IFT27 Links the BBSome to IFT for Maintenance of the Ciliary Signaling Compartment’, Developmental Cell, 31(3), pp. 279–290. doi:10.1016/j.devcel.2014.09.011.

Eliáš, M. et al. (2016) ‘A paneukaryotic genomic analysis of the small GTPase RABL2 underscores the significance of recurrent gene loss in eukaryote evolution’, Biology Direct, 11(1), p. 5. doi:10.1186/s13062-016-0107-8.

Engel, B.D., Ludington, W.B. and Marshall, W.F. (2009) ‘Intraflagellar transport particle size scales inversely with flagellar length: revisiting the balance-point length control model’, Journal of Cell Biology, 187(1), pp. 81–89. doi:10.1083/jcb.200812084.

Evans, R. et al. (2022) ‘Protein complex prediction with AlphaFold-Multimer’. bioRxiv, p. 2021.10.04.463034. doi:10.1101/2021.10.04.463034.

Funabashi, T. et al. (2018) ‘Interaction of heterotrimeric kinesin-II with IFT-B–connecting tetramer is crucial for ciliogenesis’, Journal of Cell Biology, 217(8), pp. 2867–2876. doi:10.1083/jcb.201801039.

Glaser, F. et al. (2003) ‘ConSurf: Identification of Functional Regions in Proteins by Surface-Mapping of Phylogenetic Information’, Bioinformatics, 19(1), pp. 163–164. doi:10.1093/bioinformatics/19.1.163.

Götze, M. et al. (2015) ‘Automated Assignment of MS/MS Cleavable Cross-Links in Protein 3D-Structure Analysis’, Journal of the American Society for Mass Spectrometry, 26(1), pp. 83–97. doi:10.1007/s13361-014-1001-1.

Guo, Z. et al. (2013) ‘Intermediates in the Guanine Nucleotide Exchange Reaction of Rab8 Protein Catalyzed by Guanine Nucleotide Exchange Factors Rabin8 and GRAB*’, Journal of Biological Chemistry, 288(45), pp. 32466–32474. doi:10.1074/jbc.M113.498329.

Hibbard, J.V.K. et al. (2021) ‘Protein turnover dynamics suggest a diffusion-to-capture mechanism for peri-basal body recruitment and retention of intraflagellar transport proteins’, Molecular Biology of the Cell, 32(12), pp. 1171–1180. doi:10.1091/mbc.E20-11-0717.

Hoek, H. van den et al. (2021) ‘In situ architecture of the ciliary base reveals the stepwise assembly of IFT trains’. bioRxiv, p. 2021.10.17.464685. doi:10.1101/2021.10.17.464685.

Hou, Y., Pazour, G.J. and Witman, G.B. (2004) ‘A Dynein Light Intermediate Chain, D1bLIC, Is Required for Retrograde Intraflagellar Transport’, Molecular Biology of the Cell, 15(10), pp. 4382–4394. doi:10.1091/mbc.e04-05-0377.

Iacobucci, C. et al. (2018) ‘A cross-linking/mass spectrometry workflow based on MS-cleavable cross-linkers and the MeroX software for studying protein structures and protein–protein interactions’, Nature Protocols, 13(12), pp. 2864–2889. doi:10.1038/s41596-018-0068-8.

Inglis, P.N., Blacque, O.E. and Leroux, M.R. (2009) ‘Chapter 14 - Functional Genomics of Intraflagellar Transport-Associated Proteins in C. elegans’, in King, S.M. and Pazour, G.J. (eds) Methods in Cell Biology. Academic Press (Methods in Cell Biology), pp. 267–304. doi:10.1016/S0091-679X(08)93014-4.

Jakobsen, L. et al. (2011) ‘Novel asymmetrically localizing components of human centrosomes identified by complementary proteomics methods’, The EMBO Journal, 30(8), pp. 1520–1535. doi:10.1038/emboj.2011.63.

Jordan, M.A. et al. (2018) ‘The cryo-EM structure of intraflagellar transport trains reveals how dynein is inactivated to ensure unidirectional anterograde movement in cilia’, Nature Cell Biology, 20(11), pp. 1250–1255. doi:10.1038/s41556-018-0213-1.

Jumper, J. et al. (2021) ‘Highly accurate protein structure prediction with AlphaFold’, Nature, 596(7873), pp. 583–589. doi:10.1038/s41586-021-03819-2.

Kanie, T. et al. (2017) ‘The CEP19-RABL2 GTPase Complex Binds IFT-B to Initiate Intraflagellar Transport at the Ciliary Base’, Developmental Cell, 42(1), pp. 22–36.e12. doi:10.1016/j.devcel.2017.05.016.

Keady, B.T. et al. (2012) ‘IFT25 Links the Signal-Dependent Movement of Hedgehog Components to Intraflagellar Transport’, Developmental Cell, 22(5), pp. 940–951. doi:10.1016/j.devcel.2012.04.009.

Knödler, A. et al. (2010) ‘Coordination of Rab8 and Rab11 in primary ciliogenesis’, Proceedings of the National Academy of Sciences, 107(14), pp. 6346–6351. doi:10.1073/pnas.1002401107.

Kozminski, K.G. et al. (1993) ‘A motility in the eukaryotic flagellum unrelated to flagellar beating.’, Proceedings of the National Academy of Sciences, 90(12), pp. 5519–5523. doi:10.1073/pnas.90.12.5519.

Kozminski, K.G., Beech, P.L. and Rosenbaum, J.L. (1995) ‘The Chlamydomonas kinesin-like protein FLA10 is involved in motility associated with the flagellar membrane.’, Journal of Cell Biology, 131(6), pp. 1517–1527. doi:10.1083/jcb.131.6.1517.

Kramer, M. et al. (2010) ‘Analysis of relative gene dosage and expression differences of the paralogs RABL2A and RABL2B by Pyrosequencing’, Gene, 455(1), pp. 1–7. doi:10.1016/j.gene.2010.01.005.

Landau, M. et al. (2005) ‘ConSurf 2005: the projection of evolutionary conservation scores of residues on protein structures’, Nucleic Acids Research, 33(suppl_2), pp. W299–W302. doi:10.1093/nar/gki370.

Lechtreck, K.-F. et al. (2009) ‘The Chlamydomonas reinhardtii BBSome is an IFT cargo required for export of specific signaling proteins from flagella’, Journal of Cell Biology, 187(7), pp. 1117–1132. doi:10.1083/jcb.200909183.

Lechtreck, K.F. et al. (2013) ‘Cycling of the signaling protein phospholipase D through cilia requires the BBSome only for the export phase’, Journal of Cell Biology, 201(2), pp. 249– 261. doi:10.1083/jcb.201207139.

Liew, G.M. et al. (2014) ‘The Intraflagellar Transport Protein IFT27 Promotes BBSome Exit from Cilia through the GTPase ARL6/BBS3’, Developmental Cell, 31(3), pp. 265–278. doi:10.1016/j.devcel.2014.09.004.

Lo, J.C.Y. et al. (2012) ‘RAB-Like 2 Has an Essential Role in Male Fertility, Sperm Intra-Flagellar Transport, and Tail Assembly’, PLOS Genetics, 8(10), p. e1002969. doi:10.1371/journal.pgen.1002969.

Ludington, W.B. et al. (2013) ‘Avalanche-like behavior in ciliary import’, Proceedings of the National Academy of Sciences, 110(10), pp. 3925–3930. doi:10.1073/pnas.1217354110.

Lv, B. et al. (2017) ‘Intraflagellar transport protein IFT52 recruits IFT46 to the basal body and flagella’, Journal of Cell Science, 130(9), pp. 1662–1674. doi:10.1242/jcs.200758.

Mijalkovic, J. et al. (2018) ‘Single-Molecule Turnarounds of Intraflagellar Transport at the C. elegans Ciliary Tip’, Cell Reports, 25(7), pp. 1701–1707.e2. doi:10.1016/j.celrep.2018.10.050.

Mishra, A.K. and Lambright, D.G. (2016) ‘Invited review: Small GTPases and their GAPs’, Biopolymers, 105(8), pp. 431–448. doi:10.1002/bip.22833.

Nishijima, Y. et al. (2017) ‘RABL2 interacts with the intraflagellar transport-B complex and CEP19 and participates in ciliary assembly’, Molecular Biology of the Cell, 28(12), pp. 1652– 1666. doi:10.1091/mbc.e17-01-0017.

Nottingham, R.M. et al. (2011) ‘RUTBC1 Protein, a Rab9A Effector That Activates GTP Hydrolysis by Rab32 and Rab33B Proteins’, Journal of Biological Chemistry, 286(38), pp. 33213–33222. doi:10.1074/jbc.M111.261115.

Pai, E.F. et al. (1990) ‘Refined crystal structure of the triphosphate conformation of H-ras p21 at 1.35 A resolution: implications for the mechanism of GTP hydrolysis.’, The EMBO Journal, 9(8), pp. 2351–2359.

Pan, X. et al. (2006) ‘TBC-domain GAPs for Rab GTPases accelerate GTP hydrolysis by a dual-finger mechanism’, Nature, 442(7100), pp. 303–306. doi:10.1038/nature04847.

Pedersen, L.B. and Rosenbaum, J.L. (2008) ‘Chapter Two Intraflagellar Transport (IFT): Role in Ciliary Assembly, Resorption and Signalling’, in Current Topics in Developmental Biology. Academic Press (Ciliary Function in Mammalian Development), pp. 23–61. doi:10.1016/S0070-2153(08)00802-8.

Pigino, G. et al. (2009) ‘Electron-tomographic analysis of intraflagellar transport particle trains in situ’, Journal of Cell Biology, 187(1), pp. 135–148. doi:10.1083/jcb.200905103.

Qin, H. et al. (2007) ‘Intraflagellar Transport Protein 27 Is a Small G Protein Involved in Cell-Cycle Control’, Current Biology, 17(3), pp. 193–202. doi:10.1016/j.cub.2006.12.040.

Quidwai, T. et al. (2021) ‘A WDR35-dependent coat protein complex transports ciliary membrane cargo vesicles to cilia’, eLife. Edited by R.A. Kahn, V. Malhotra, and R.A. Kahn, 10, p. e69786. doi:10.7554/eLife.69786.

Rosenbaum, J.L. and Witman, G.B. (2002) ‘Intraflagellar transport’, Nature Reviews Molecular Cell Biology, 3(11), pp. 813–825. doi:10.1038/nrm952.

Schafer, J.C. et al. (2006) ‘IFTA-2 is a conserved cilia protein involved in pathways regulating longevity and dauer formation in Caenorhabditis elegans’, Journal of Cell Science, 119(Pt 19), pp. 4088–4100. doi:10.1242/jcs.03187.

Scheffzek, K. and Ahmadian, M.R. (2005) ‘GTPase activating proteins: structural and functional insights 18 years after discovery’, Cellular and Molecular Life Sciences CMLS, 62(24), pp. 3014–3038. doi:10.1007/s00018-005-5136-x.

Scheffzek, K. and Shivalingaiah, G. (2019) ‘Ras-Specific GTPase-Activating Proteins—Structures, Mechanisms, and Interactions’, Cold Spring Harbor Perspectives in Medicine, 9(3), p. a031500. doi:10.1101/cshperspect.a031500.

Seewald, M.J. et al. (2002) ‘RanGAP mediates GTP hydrolysis without an arginine finger’, Nature, 415(6872), pp. 662–666. doi:10.1038/415662a.

Shalata, A. et al. (2013) ‘Morbid Obesity Resulting from Inactivation of the Ciliary Protein CEP19 in Humans and Mice’, The American Journal of Human Genetics, 93(6), pp. 1061– 1071. doi:10.1016/j.ajhg.2013.10.025.

Silva, D.A. et al. (2012) ‘The RABL5 homolog IFT22 regulates the cellular pool size and the amount of IFT particles partitioned to the flagellar compartment in Chlamydomonas reinhardtii’, Cytoskeleton, 69(1), pp. 33–48. doi:10.1002/cm.20546.

Sorokin, S. (1962) ‘CENTRIOLES AND THE FORMATION OF RUDIMENTARY CILIA BY FIBROBLASTS AND SMOOTH MUSCLE CELLS’, Journal of Cell Biology, 15(2), pp. 363–377. doi:10.1083/jcb.15.2.363.

Stepanek, L. and Pigino, G. (2016) ‘Microtubule doublets are double-track railways for intraflagellar transport trains’, Science, 352(6286), pp. 721–724. doi:10.1126/science.aaf4594.

Taschner, M. et al. (2014) ‘Crystal structures of IFT70/52 and IFT52/46 provide insight into intraflagellar transport B core complex assembly’, Journal of Cell Biology, 207(2), pp. 269– 282. doi:10.1083/jcb.201408002.

Taschner, M. et al. (2016) ‘Intraflagellar transport proteins 172, 80, 57, 54, 38, and 20 form a stable tubulin-binding IFT-B2 complex’, The EMBO Journal, 35(7), pp. 773–790. doi:10.15252/embj.201593164.

Taschner, M. and Lorentzen, E. (2016a) ‘The Intraflagellar Transport Machinery’, Cold Spring Harbor Perspectives in Biology, 8(10), p. a028092. doi:10.1101/cshperspect.a028092.

Taschner, M. and Lorentzen, E. (2016b) ‘The Intraflagellar Transport Machinery’, Cold Spring Harbor Perspectives in Biology, 8(10), p. a028092. doi:10.1101/cshperspect.a028092.

Vetter, M. et al. (2015a) ‘Structure of Rab11–FIP3–Rabin8 reveals simultaneous binding of FIP3 and Rabin8 effectors to Rab11’, Nature Structural & Molecular Biology, 22(9), pp. 695–702. doi:10.1038/nsmb.3065.

Vetter, M. et al. (2015b) ‘Structure of Rab11–FIP3–Rabin8 reveals simultaneous binding of FIP3 and Rabin8 effectors to Rab11’, Nature Structural & Molecular Biology, 22(9), pp. 695–702. doi:10.1038/nsmb.3065.

Wachter, S. et al. (2019) ‘Binding of IFT22 to the intraflagellar transport complex is essential for flagellum assembly’, The EMBO Journal, 38(9), p. e101251. doi:10.15252/embj.2018101251.

Wang, Z. et al. (2009) ‘Intraflagellar Transport (IFT) Protein IFT25 Is a Phosphoprotein Component of IFT Complex B and Physically Interacts with IFT27 in Chlamydomonas’, PLOS ONE, 4(5), p. e5384. doi:10.1371/journal.pone.0005384.

Wingfield, J.L. et al. (2017) ‘IFT trains in different stages of assembly queue at the ciliary base for consecutive release into the cilium’, eLife. Edited by E.J.G. Peterman, 6, p. e26609. doi:10.7554/eLife.26609.

Wingfield, J.L. et al. (2021) ‘In vivo imaging shows continued association of several IFT-A, IFT-B and dynein complexes while IFT trains U-turn at the tip’, Journal of Cell Science, 134(18), p. jcs259010. doi:10.1242/jcs.259010.

Wittinghofer, A. and Vetter, I.R. (2011) ‘Structure-function relationships of the G domain, a canonical switch motif’, Annual Review of Biochemistry, 80, pp. 943–971. doi:10.1146/annurev-biochem-062708-134043.

Wong, A.C.C. et al. (1999) ‘Two Novel Human RAB Genes with Near Identical Sequence Each Map to a Telomere-Associated Region: The Subtelomeric Region of 22q13.3 and the Ancestral Telomere Band 2q13’, Genomics, 59(3), pp. 326–334. doi:10.1006/geno.1999.5889.

Xue, B. et al. (2020) ‘Intraflagellar transport protein RABL5/IFT22 recruits the BBSome to the basal body through the GTPase ARL6/BBS3’, Proceedings of the National Academy of Sciences, 117(5), pp. 2496–2505. doi:10.1073/pnas.1901665117.

Zhou, Z. et al. (2022) ‘Impaired cooperation between IFT74/BBS22–IFT81 and IFT25– IFT27/BBS19 causes Bardet-Biedl syndrome’, Human Molecular Genetics, 31(10), pp. 1681–1693. doi:10.1093/hmg/ddab354.

